# WGT-aware analysis reveals increased complexity in the yohimbane biosynthesis pathway of *Rauvolfia tetraphylla*

**DOI:** 10.64898/2026.02.22.707333

**Authors:** Mratunjay Dwivedi, Nagarjun Vijay

## Abstract

Arising from Stander et al. Communications Biology https://doi.org/10.1038/s42003-023-05574-8 (2023)

*Rauvolfia tetraphylla* (Apocynaceae), distributed across Mexico, Tropical America, and Southeast Asia, is known to produce pharmacologically important monoterpene indole alkaloids such as ajmaline, reserpiline, yohimbine, and heteroyohimbanes^1^. Elucidating the enzymatic pathways^2^ underlying their biosynthesis is therefore essential for enhancing our understanding of MIA metabolism and enabling their industrial production. In a recent edition of Communication Biology, Stander et al. [STAN23]^3^ presented a high-quality assembly for *R. tetraphylla* and uncovered the yohimbane biosynthesis pathway. Based on their proteomics and transcriptomic analysis, two major findings emerged: (1) a medium chain dehydrogenase/reductase (MDR), yohimbane synthase (YOS), produces four yohimbane diastereomers, and (2) three MDR transcripts (MDRT), MSTRG.5530, MSTRG.5531, and MSTRG.5534, along with geissoschizine synthase (GS, MSTRG.5528), can generate a mixture of yohimbane isomers. However, our reanalysis of the raw sequencing reads incorporating haplotype phasing and expression profiling revealed that most candidate genes exist as three homeologous copies with distinct tissue-specific expression. Furthermore, we demonstrate preferential interactions among specific MDRT and GS homeologs, indicating complex cross-subchromosomal enzymatic organisation overlooked in STAN23. Furthermore, several reported candidate genes are chimeric, indicating that incorrect estimation of whole-genome triplication and lack of haplotype phasing can severely limit robust identification and functional characterisation of the yohimbane biosynthetic pathway.

## Introduction

STAN23 generated a high-quality de novo genome assembly of *R. tetraphylla*, using long ONT reads and Illumina-based polishing and also generated extensive transcriptomic data across various tissue types. Their study constitutes the first characterisation of yohimbane biosynthetic mechanisms; integrating gene co-expression analysis, deep learning classification, in vitro biochemical assays and LC-MS analysis, they identified multiple enzymatic routes to yohimbane-type MIAs, including a multifunctional MDR; YOS, as well as three distinct MDR genes which couple with GS in a two-enzyme pathway to produce yohimbane and its stereoisomers using strictosidine aglycone as a precursor. However, the well-foundedness of the results is limited by a methodological oversight: the study does not take into account an independent whole-genome triplication event (WGT) in *R. tetraphylla* other than the one shared across Eudicotyledons, thus limiting the candidates identified in yohimbane biosynthesis and undermining the complexity of the biosynthetic modalities^4,5^.

In another published study, Lezin et al. [LEZ24]^6^ presented an independent assembly of *R. tetraphylla*, which incorporates an independent ancestral WGT compared to other *Apocynaceae* species, revealing a more complex genome architecture with three homeologous copies for each chromosome. Interestingly, both assemblies were generated from the same raw nanopore sequencing data and employed similar analytical approaches, such as Ks distribution curves (STAN23 Fig. 2c; LEZ24 Fig. 1b), yet yielded contrasting estimates of the WGT event. The smudge plots^7^ generated from raw reads revealed distinct genomic architecture between *R. tetraphylla* and its congeneric species, *Rauvolfia serpentina*^8^ (**Fig. 1A, 1B**). While *R. serpentina* exhibited clear, discrete peaks consistent with a diploid genome, *R. tetraphylla* displayed a broader peak pattern, indicating a higher ploidy level. To further resolve such ambiguities in the WGT status and consequently the genomic organisation of *R. tetraphylla*, we employed a haplotype phasing approach on chromosome 11 of the WGT-aware LEZ24 assembly by identifying haplotype-defining polymorphisms (HDPs) that can distinguish among subchromosomes (haplotypes) 11a, 11b, and 11c^9^. We demonstrated alignment of distinct long reads to their respective haplotypes based on the allelic states at those HDPs. Quantification of this read support validated the phasing correctness (**Fig. 1C, 1D**; Supplementary Text S1, S2 and Supplementary Figs. S1-S4). In summary, our analysis provided strong support for an additional WGT event and confirmed the structural integrity of the LEZ24 genome assembly.

**Fig. 1:**
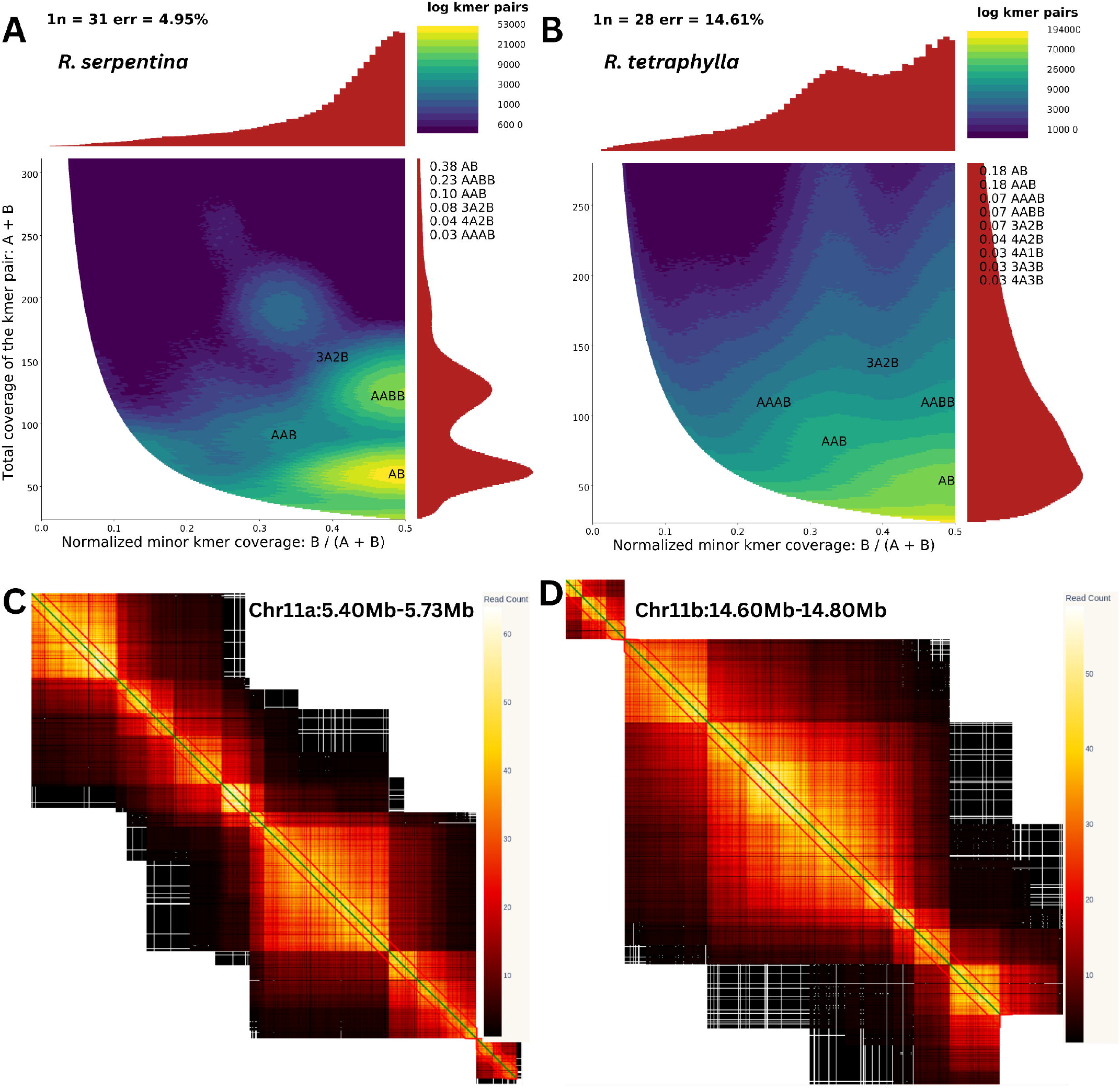
Ploidy assessment and mapping-based haplotype phasing validation of *R. tetraphylla* LEZ24 genome. **A** Smudgeplot for *R. serpentina*; the two clear peaks indicate a diploid genome. **B** Smudgeplot for *R. tetraphylla*, the broad peak indicates a higher-ploidy genome. **C** Haplotype linkage heatmaps for HDP pairs for chromosome 11a using Rte_Chr11a as reference for mapping. Each square block represents a HDP pair. The colour scale on the right indicates the strength of read support for the 11a allele for each HDP pair, with yellow denoting the highest and black the lowest. A strong diagonal pattern reflects linkage between adjacent HDPs, indicating haplotype continuity; read support diminishes as the distance between HDP pairs increases. Regions with sparse HDPs exhibit weak or fragmented diagonal support. Red outlines highlight strongly phased haplotype blocks. **D** Haplotype linkage heatmap for chromosome 11b using Rte_Chr11b as reference; visualising read support for 11b allele.

**Fig. 2:**
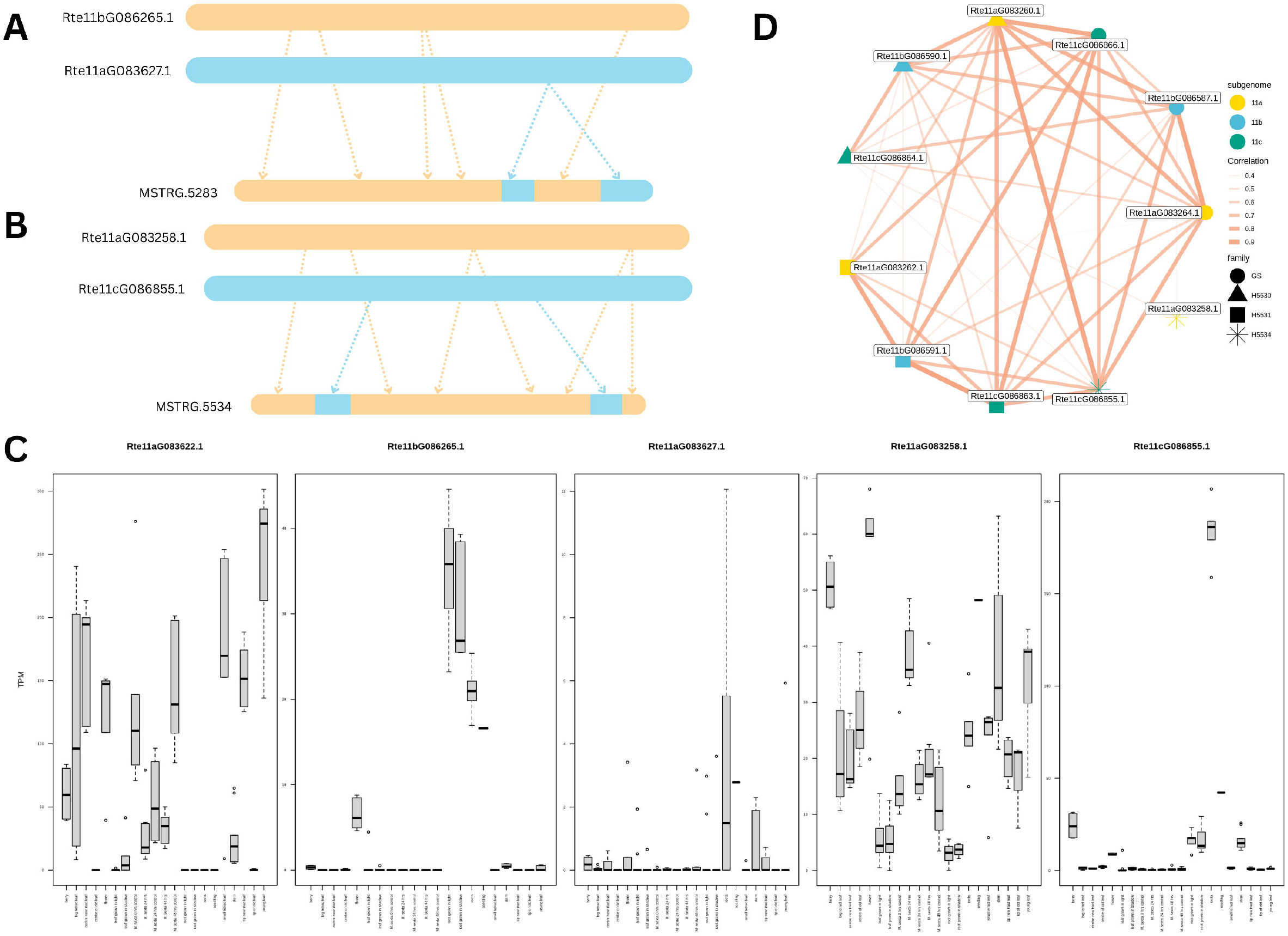
Mosaic transcript structure, coexpression and expression divergence across yohimbane candidate gene homeologs. **A** Schematic representation of the mosaic structure of MSTRG.5283 representing contributions from its homeologous 11b (Rte11bG086265.1) and 11a (Rte11aG083627.1) copy. Single colour blocks represent a pure haplotype. **B** Schematic representation of the mosaic structure of MSTRG.5534 representing contributions from homeologous 11a (Rte11aG083258.1) and 11c (Rte11cG086855.1) copy. **C** Tissue and condition-specific expression profiles of representative homeologous genes are shown in panels (A, B). Boxplots summarise expression across biological replicates for each tissue or condition, illustrating differential expression patterns and functional divergence among homeologs. **D** Co-expression network of yohimbane candidate genes in the two-enzyme pathway. Nodes represent genes and are colored by the subgenome of origin (11a, 11b, 11c). Node shapes denote gene family membership (GS, HSS30, HSS31, HSS34). Edges represent pairwise expression correlations, with line thickness and opacity proportional to correlation strength.

## Results

Reanalysis of yohimbane biosynthetic gene candidates using the LEZ24 genome, employing BLAST^10^, OrthoFinder^11^, co-expression analysis, deep learning classification and domain characterisation (Supplementary Text S3), we found that most candidate genes, including YOS (MSTRG.5283), GS and MDRs (MSTRG.5530, MSTRG.5531, MSTRG.5534) exist as three homeologous copies distributed across subchromosomes 11a, 11b, and 11c. Specifically, GS corresponds to Rte11aG083264.1, Rte11bG086587.1, and Rte11cG086866.1; MSTRG.5530 to Rte11aG083260.1, Rte11bG086590.1, and Rte11cG086864.1; MSTRG.5531 to Rte11aG083262.1, Rte11bG086591.1, and Rte11cG086863.1; whereas MSTRG.5534 is detected only on 11a (Rte11aG083258.1) and 11c (Rte11cG086855.1). Additionally, YOS is present as Rte11aG083627.1 and Rte11bG086265.1, alongside a highly identical paralogous copy, Rte11aG083622.1 (Supplementary Data S1). Three homeologous copies are expected per gene across the subchromosomes due to the WGT event; however, certain copies could not be identified, suggesting putative divergence or gene loss.

Since haplotypes are collapsed in the STAN23 assembly, we investigated which one of the three homeologous copies in the LEZ24 genome corresponds to the STAN23 genes. This was achieved by aligning all homeologous copies of each gene and designating mismatching non-gap positions as HDPs (Supplementary Text S4). Our analysis showed that MSTRG.5528, 5530, and 5531 predominantly align with their corresponding 11c homeolog, exhibiting high allelic similarity at HDPs. In contrast, MSTRG.5534 shows strong allelic support for the 11a homeolog but also considerable similarity to the 11c copy at certain HDPs. Similarly, YOS primarily aligns with its 11b homeolog, yet also displays substantial support for the 11a copy alleles at certain HDPs, indicating that a few genes used in the STAN23 study represent a chimeric sequence (**Fig. 2A, 2B**). These mismatches include both synonymous and non-synonymous variants that may cause amino acid changes within the functional protein domains. This suggests that the functional assays based on MSTRG.5534 and MSTRG.5283 may not accurately reflect the true biological activity, as the chimeric enzymes could differ in protein function or expression compared to their native LEZ24 counterparts.

Expression profiling via RNA-seq revealed that the single-copy transcripts used by STAN23 for functional validation do not correspond to the most highly expressed homeologs (Supplementary Data S2, S3). For instance, GS shows the highest expression in its 11b copy; MSTRG.5530 in its 11a copy; MSTRG.5531 and MSTRG.5534 in their 11b and 11c homeolog respectively (Supplementary Figs. S5-S7). Similarly, for YOS (MSTRG.5283), the 20 amino-acid longer Rte11aG083622.1 transcript exhibits substantially higher expression (**Fig. 2C**) than both Rte11aG083627.1 and Rte11bG086265.1, underscoring the dominance of different gene copies than that identified by STAN23. This implies that the yohimbane biosynthesis in *R. tetraphylla* is governed by homeolog-specific expression, and functional studies involving multiple homeologs are essential to accurately resolve its enzymatic pathways.

The expression patterns of gene copies also exhibit strong variation across tissues, suggesting subfunctionalisation and neofunctionalisation with distinct homeologs being primarily utilised among tissues for yohimbane biosynthesis. We hypothesise that, within the two-enzyme pathway, specific copies of MSTRG.5530, 5531, and 5534 preferentially interact with particular GS homeologs to produce yohimbane. To test this hypothesis, we performed co-expression network analysis using the LEZ24 assembly. This approach revealed that all the MSTRG.5531 homeologs exhibit a strong correlation with GS-11c copy. Additionally, MSTRG.5530 homeologs display cross-subchromosomal correlations with GS homeologs; the 11c copy correlates with GS-11b copy, while the 11b and 11a copies show strong correlation with GS-11c copy (**Fig. 2D**). These findings demonstrate that the distinct copies of MSTRG.5530, 5531, and 5534 preferentially associate with specific GS homeologs, revealing complex cross-subchromosomal co-expression patterns overlooked in the previous STAN23 analyses. Consequently, the gene pairs utilised in STAN23 functional assays do not correspond to the primary homeolog pairs active in planta, thus limiting their biological relevance.

## Discussion

In conclusion, the STAN23 study is fundamentally limited due to an inaccurate estimation of the WGT event, which led to the omission of additional candidate gene homeologs with tissue-specific expression, consequently overlooking potential subfunctionalisation and neofunctionalization across the gene copies. The study also fails to capture critical interactions within the two-enzyme pathway involving GS and associated MDRT copies, thereby undermining the complexity of yohimbane biosynthesis. The absence of haplotype phasing further contributed to the identification and downstream analysis of chimeric candidate genes, which may not accurately represent the native biological sequences or their functional characteristics. The correct gene sequences will not only better reflect the endogenous biological activity but may also enhance enzymatic efficiency, thus serving as superior candidates for industrial production. We emphasise that ploidy assessment using raw sequencing reads combined with mapping-based haplotype phasing is essential as a post-assembly validation step to prevent misinterpretations caused by assembly artifacts^12^. This approach is particularly crucial when assembled gene sequences are intended for subsequent functional studies. Future research should prioritise the yohimbane candidate gene homeologs identified through the LEZ24 assembly. Additionally, co-expressed GS-MDRT gene pairs revealed in this analysis should be the focus of functional characterisation to elucidate their specific functions in yohimbane biosynthesis and accurately reflect the key enzymatic contributors in the plant.

## Methods

In our analysis, we performed mapping-based haplotype phasing of *R. tetraphylla* chromosome 11 by identifying HDPs through pairwise alignments of the three subchromosomes (11a, 11b, and 11c). Nanopore reads were then categorised based on allelic states at these HDPs, and read support for consistent alleles was quantified to validate the haplotype correctness. This approach substantiated the presence of an additional WGT event and confirmed the accuracy of the LEZ24 genome assembly.

Expression abundance was quantified as transcripts per million (TPM) by pseudo-aligning RNA-seq reads from 150 samples using Salmon^13^. Transcripts with TPM > 10 in at least six samples were retained for downstream analysis. Expression robustness was validated using cross-expression analysis (Supplementary Text S5, Supplementary Figs. S8-S10). Co-expression networks integrating gene expression were generated using Spearman’s rank correlation (Supplementary Text S6, S7 and Supplementary Figs. S11-S27) ranked by highest reciprocal ranks (HRR); known MIA genes were used as baits to identify novel co-expressed candidates, with network visualisation performed using igraph (v2.2.1)^14^ in R (v4.3). A focused co-expression network was further generated using only yohimbane candidate gene homeologs to investigate their specific interactions. Correlation robustness against tissue composition was evaluated using leave-one-tissue-out (LOTO) analysis (Supplementary Text S8, Supplementary Figs. S28-S32). Deep learning classification of MIA-related genes was performed using a feedforward artificial neural network trained with the H2O library (v3.46) in R (v4.3), employing a labelled dataset of 96 MIA and 28,087 non-MIA transcripts derived from BUSCO^15^ orthologs (Supplementary Text S9 and Supplementary Data S4).

## Author contributions

Mratunjay Dwivedi: Conceptualisation, Formal analysis, Investigation, Visualisation, Validation, Writing - original draft, Writing - review & editing. Nagarjun Vijay: Conceptualization, Resources, Writing - original draft, Writing - review & editing, Funding acquisition, Project administration, Supervision.

## Data availability

Data are available at https://github.com/MrityunjayCEGLAB/MatterArising_comm_biol

## Acknowledgements

We thank IISER Bhopal for supporting MD with a PhD scholarship. The Department of Biotechnology, Ministry of Science and Technology, India (Grant no. BT/11/IYBA/2018/03) and the Science and Engineering Research Board (Grant no. ECR/2017/001430) provided funds for computational resources (i.e., Har Gobind Khorana Computational Biology cluster) used.

