## Supplementary Materials for "WGT-aware analysis reveals increased complexity in the yohimbane biosynthesis pathway of *Rauvolfia tetraphylla*"

Arising from Stander et al. Communications Biology https://doi.org/10.1038/s42003-023-05574-8 (2023)

#### 1. Mapping-based haplotype phasing validation for the LEZ24 genome

In order to validate haplotype phasing in the LEZ24[1] genome, we employed the mapping-based phasing approach. Although this method can be used to validate any chromosome copies, we exclusively validated the Rte_Chr11a, Rte_Chr11b, and Rte_Chr11c, because our candidate genes and other ADHs were present on Chromosome 11. In this approach, pairwise alignments were performed by mapping Rte_Chr11b and Rte_Chr11c independently onto Rte_Chr11a. In order to minimise redundancy arising from secondary or supplementary mappings, the resulting BAM files were filtered to retain non-overlapping alignments per reference position, such that each reference base was represented by at most one alignment. Filtered BAM files were parsed using custom python scripts to extract base-level alignment information. For each alignment, the mapped base at every aligned reference position was compared to the corresponding reference base in Rte_Chr11a. Positions at which the mapped base differed from the reference base were recorded as mismatches in tab-separated files for both Rte_Chr11b-to-Rte_Chr11a and Rte_Chr11c-to-Rte_Chr11a mappings.

Reference positions shared between the two alignments (i.e., positions in Rte_Chr11a that were covered by both Rte_Chr11b and Rte_Chr11c alignments) were retained. Among these shared positions, sites exhibiting base differences among the three chromosomes were designated as haplotype-defining polymorphisms (HDPs), distinguishing Rte_Chr11a, Rte_Chr11b, and Rte_Chr11c. To assign long reads to specific haplotypes, Oxford Nanopore reads mapped to chromosome 11 were parsed to extract the bases present at the identified HDP positions. For each read, the observed bases at HDPs were compared against the expected alleles for each chromosome, and allelic support for Rte_Chr11a, Rte_Chr11b or Rte_Chr11c was quantified. For robustness and sensitivity, reads were assigned to a specific homeologous chromosome if at least 70% of the HDPs covered by that read supported a single chromosome. Finally, allelic support for pairs of HDP positions, by the same read, was quantified across haplotypes. These data were used to assess relative read support and continuity among chromosome-11 haplotypes **(Supplementary Figure S1)**.

#### 2. Whole chromosomal alignments among chromosome 11 subchromosomes

To qualitatively assess sequence conservation and collinearity among the three chromosome-11 subchromosomes, Rte_Chr11a, Rte_Chr11b, and Rte_Chr11c, we performed whole-chromosome alignments using progressiveMauve **(Supplementary Figure S2-S4)**. Pairwise alignments were generated for each subchromosome combination, 11a–11b, 11a–11c and 11b–11c using default progressiveMauve settings. The resulting alignments were visualised using the pMauve viewer. Conserved segments were interpreted as locally collinear blocks (LCBs), representing regions of homologous sequence shared between the two input chromosomes. LCBs are displayed as colored blocks; blocks of the same colour correspond to the same homologous segment across the aligned sequences. Connecting lines denote aligned regions between the two chromosomes, where forward-orientation links indicate collinear alignment and inverted links indicate alignment in reverse orientation (consistent with inversions).

Across comparisons, we observed clear conservation of multiple long LCBs, supporting broad homology among the three subchromosomes and confirming their derivation from a shared ancestral chromosome segment. These results are directly relevant to mapping-based haplotype phasing. First, the presence of long collinear blocks supports the validity of identifying haplotype-defining polymorphisms (HDPs) via mapping-based pairwise alignments, as homologous regions are sufficiently conserved to permit high-confidence base comparisons.

#### 3. Blast and OrthoFinder detection of candidate gene copies

To re-evaluate candidate yohimbane biosynthetic genes identified by STAN23[2] within the WGT-aware LEZ24 genome assembly[3], we analysed the corresponding MDR copies of MSTRG.5283 (YOS), MSTRG.5528 (GS), MSTRG.5530, MSTRG.5531, and MSTRG.5534. Each transcript was used as a query in BLAST[4] searches against the LEZ24 genome assembly. Hits were stringently filtered to retain only high-confidence matches exhibiting ≥95% query coverage and ≥80% sequence identity, ensuring reliable homeolog identification **(Supplementary Data S1)**. Absence of certain homeologous copies likely reflects gene loss or divergence events.

These BLAST results were further supported by Orthofinder[5] analyses, which clustered corresponding copies into orthogroups when run with the LEZ24 genome alongside outgroup species such as Catharanthus roseus. This integrated approach validated the presence and evolutionary relationships of homeologous gene copies distributed across the subchromosomes. To assess functional conservation among the identified homeologs, Pfam[6] domain analysis was performed for each candidate gene and protein model across the three subchromosomes. The conserved Pfam domain architectures observed within YOS, GS, and MDR candidate sets reinforce their functional homology and substantiate the inferred homeologous relationships.

#### 4. Multiple sequence alignment–based detection of mosaic (chimeric) transcripts

To assess whether STAN23 transcripts represent single-copy homeologs or chimeric sequences derived from multiple homeologous copies of the LEZ24 genome, we performed a multiple sequence alignment (MSA) analysis using Clustal Omega[7]. For each candidate, transcripts MSTRG.5283, MSTRG.5528, MSTRG.5530, MSTRG.5531 and MSTRG.5534, the transcript sequence was aligned together with their corresponding homeologous gene copies from the LEZ24 genome. Positions exhibiting base differences among these sequences were designated as HDPs and used for differentiating these highly similar sequences. For each transcript, the allelic state at each HDP was compared to the expected base in the 11a, 11b or 11c copies to determine whether the transcript consistently matched a single homeolog or exhibited allelic support for multiple homeologs at HDPs, indicative of a mosaic sequence.

We repeated this analysis using the PCR-amplified Sanger-sequenced loci OR514630.1 and OR514626.1, corresponding to MSTRG.5534 and MSTRG.5283, respectively, and identified chimeric signatures in the corresponding genes based on RNA-seq data. Additionally, we analysed the primers (STAN23 Supplementary Table S6) used for MSTRG.5283 and MSTRG.5534; BLAST results indicate that these primers preferentially amplify a single WGT-derived homeolog, specifically the 11a copy in both cases, which does not correspond to the most highly expressed homeolog among the detected 11a, 11b, and 11c copies. This can be a result of artefactual recombination due to PCR amplification, especially in polyploid genomes where highly similar templates coexist. PCR amplification can introduce artefactual recombination, especially in polyploid genomes where highly similar templates coexist. Standard PCR protocols designed for diploid organisms may produce chimeric amplicons due to template switching or incomplete extension events when applied to polyploid species[8,9].

#### 5. <u>Crossmapping analysis</u>

To enhance the robustness of our analysis, we tested whether or not reads originating from one homeolog do not map to its another homeolog and thus lead to non-biological expression profiles. For this, we generated artificial reads (20x for whole-genome and 100x for GS, MSTRG.5530, MSTRG.5531, and MSTRG.5534 homeologs) from art-ngs[10] simulating the Hiseq2500-like sequencing instrument, performed its expression profiling using salmon and visualised whether the reads known to originate from a certain transcript shows cross expression with other transcripts or not.

We plotted cross-expression using LEZ24 transcripts along with stringtie[11] generated transcripts **(Supplementary Figure S8)**, using only LEZ24 transcripts **(Supplementary Figure S9)** and a focused plot for GS, MSTRG.5530, MSTRG.5531, and MSTRG.5534 homeologs **(Supplementary Figure S10)**. We observed that stringtie-generated transcripts show a lot of crossmapping, indicating many of them are duplicates or highly similar to each other. While the LEZ24 annotated transcripts show negligible crossmapping except for the isoforms of those transcripts, indicating that the expression values among homeolog is correct and the expression values can be confidently interpreted.

#### 6. <u>Normality analysis for RNA-seq data</u>

To determine the appropriate correlation tests between GS and MSTRG.5530, MSTRG.5531, and MSTRG.5534 homeologs, we first assessed the distribution of gene expression (TPM values) across samples. Gene-wise normality tests were conducted on the TPM expression matrix; expression values were parsed into a numeric matrix with transcripts as rows and RNA-seq samples as columns. For each transcript with at least three non-zero TPM values, we extracted the expression vector across all samples and assessed normality using the Shapiro–Wilk test. The null hypothesis states that the data is normally distributed. For transcripts with p-values less than 0.05, we reject the null hypothesis. We observed that most of the transcripts **(Supplementary Figure S11, S12)** showed p-value < 0.05, indicating that the transcript expression vectors follow a non-normal distribution. Therefore, non-parametric tests such as Spearman's rank correlation, which do not require a normal sample distribution, would be more suitable than Pearson correlation for coexpression analysis between GS-MDR pairs.

To evaluate whether the specific GS-MDR transcript expression values followed a normal distribution, we performed normality diagnostics using quantile–quantile (QQ) plots **(Supplementary Figure S13-S23)**. TPM values across all samples were compared to the expected quantiles of a theoretical normal distribution. Deviations from the reference diagonal line representing normally distributed points were interpreted as deviations from normality. We observed zero inflation and heavy-tailed distribution at high quantiles representing non-normal distribution.

#### 7. Coexpression analysis

Coexpression network analysis was performed by constructing a matrix of gene/transcript profiles derived from paired samples. Transcripts with TPM value exceeding 10 in more than six samples were filtered for the analysis. For each gene-gene pair within the transcriptome, the pairwise Pearson's correlation coefficient was computed and ranked according to the highest reciprocal ranks (HRR), consistent with the STAN23 analysis. Genes exhibiting the strongest co-expression with each MIA-like homolog were identified using an HRR threshold of less than 30. The relationships among co-expressed genes were visualised using the igraph (v2.2.1)[12] package in R. Known MIA genes from other species served as bait for the global coexpression network, with coexpressed neighbours representing potentially novel MIA genes **(Supplementary Figure S24)**. A more focused coexpression analysis was performed using only GS-MDR, employing Spearman rank correlation due to the non-normal distribution of its gene expression vector **(Supplementary Figures S25–S27).**

#### 8. LOTO (Leave One Tissue Out) Correlation stability

Different MDR homeolog copies (MSTRG.5530, MSTRG.5531, and MSTRG.5534) show correlation with specific GS homeologs. To ensure the robustness of this analysis, we validated whether these correlations are driven by specific tissues or reflect overall co-expression. This was done by sequentially excluding all samples from each tissue type and recalculating the correlation between MDRT homeologs and the GS copy. The correlations remained consistent throughout, and excluding any tissue type did not significantly affect the correlation values. This confirms that the observed correlations are biologically relevant and not overestimated due to any particular tissue **(Supplementary Figures S28–S33)**.

#### 9. Deep learning

To predict genes associated with the MIA biosynthetic pathway from the gene expression atlas, we developed a supervised deep-learning classifier based on transcript expression profiles. Transcript levels were obtained from the filtered TPM expression matrix, with transcripts organised as rows and samples as columns. A curated positive set of known MIA-related transcripts was labelled as true positives ("MIA"), while transcripts corresponding to conserved BUSCO[13] genes—presumed housekeeping genes not involved in specialised MIA biosynthesis—were labeled as true negatives ("nonMIA") **(Supplementary Data S4)**. All other transcripts were categorized as unlabeled ("others") and retained for genome-wide prediction but excluded from model training.

Model training and prediction were performed using the H2O library (v3.46.0.8) in R (4.4.3). The labelled data were split into training and validation subsets with an 80/20 ratio using a fixed seed of 42. To address class imbalance between MIA and nonMIA categories, class balancing was enabled during training. We trained a series of feedforward neural network models using backpropagation for 100 epochs with 10-fold cross-validation. These models included single hidden-layer networks with 8, 16, and 64 neurons, as well as larger networks (200–500 neurons) trained with L2 regularisation (l2 = 1e−5) and an input dropout ratio of 0.2. Reproducibility was ensured by fixing random seeds. Following training, each model was applied to the complete expression atlas, including unlabeled transcripts, to generate genome-wide MIA predictions. Several WGT-derived gene homeologs of MSTRG.5283, MSTRG.5528, MSTRG.5530, MSTRG.5531, and MSTRG.5534 were predicted. Consequently, many of the predicted MIA genes represent strong candidates which can be used for functional analysis to identify additional genes involved in MIA biosynthesis.

### Supplementary Figures

#### WGT-aware analysis reveals increased complexity in the yohimbane biosynthesis pathway of *Rauvolfia tetraphylla*

Mratunjay Dwivedi, Nagarjun Vijay

Computational Evolutionary Genomics Lab, Department of Biological Sciences, IISER Bhopal, Bhauri, Madhya Pradesh, India

Arising from Stander et al. Communications Biology https://doi.org/10.1038/s42003-023-05574-8 (2023)

**Chr11c:12.2Mb-12.8Mb**

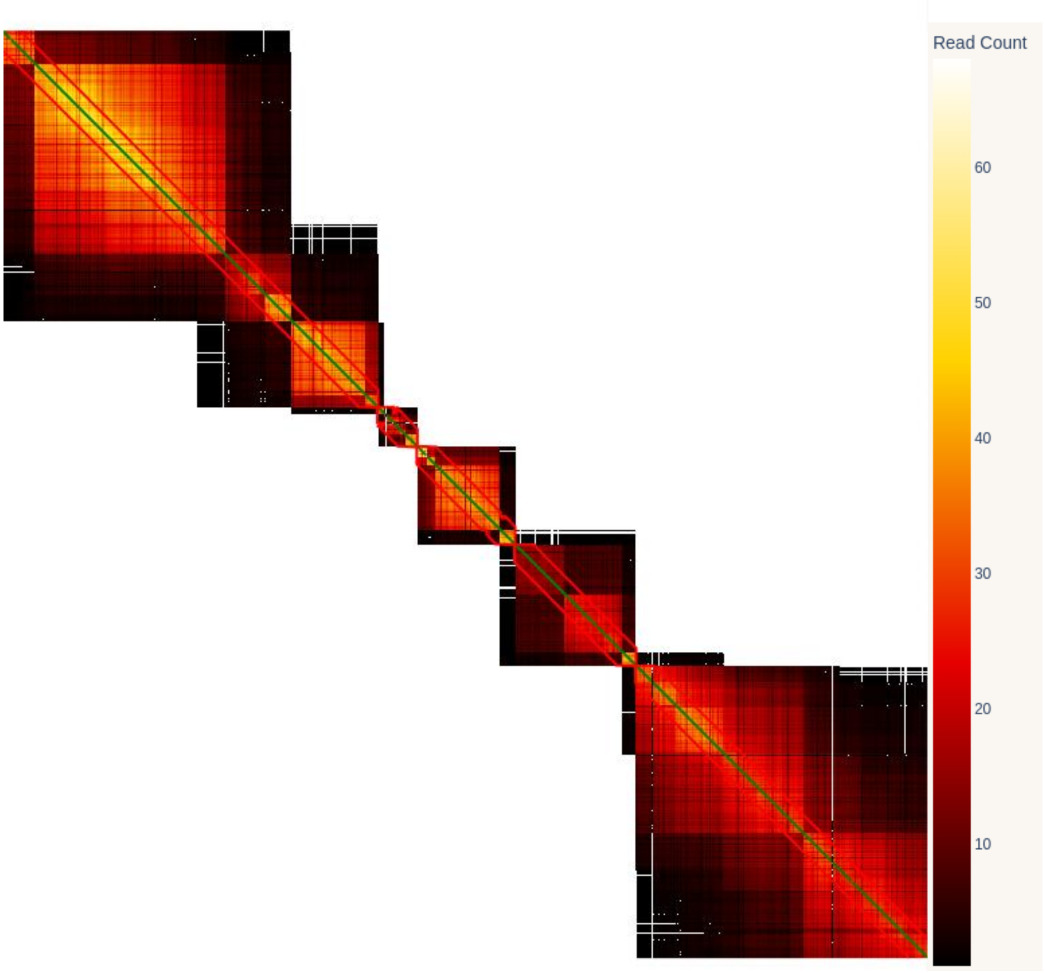


**Supplementary Figure S1: Haplotype linkage heatmaps for HDP pairs using Rte_Chr11c as reference:** Each square block represents a HDP pair. The color scale on the right indicates the strength of read support for 11a allele for each HDP pair, with yellow denoting the highest and black the lowest. A strong diagonal pattern reflects linkage between adjacent HDPs, indicating haplotype continuity; read support diminishes as the distance between HDP pairs increases. Regions with sparse HDPs exhibit weak or fragmented diagonal support. Red outlines highlight strongly phased haplotype blocks.

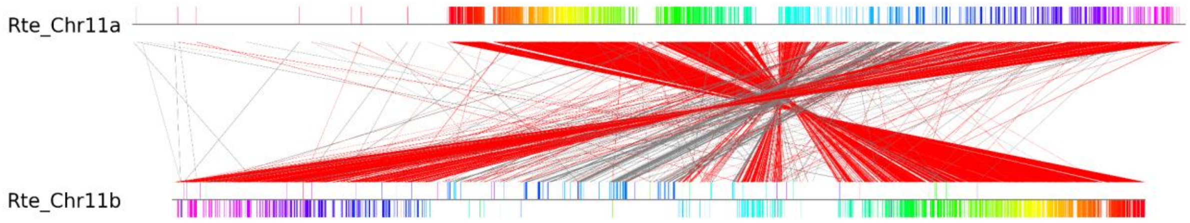


**Supplementary Figure S2: Synteny of 11a and 11b chromosomes in LEZ24 genome:** This figure represents a synteny plot generated by Progressive Mauve between Rte_Chr11a and Rte_Chr11b chromosomes. Colored blocks indicate locally collinear blocks (LCBs) across the two chromosomes. The intersecting lines between them denote the LCB strikethrough lines.

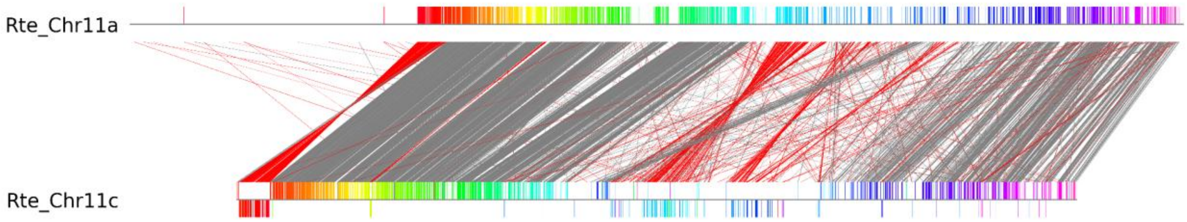


**Supplementary Figure S3: Synteny of 11a and 11c chromosomes in LEZ24 genome:** This figure represents a synteny plot generated by Progressive Mauve between Rte_Chr11a and Rte_Chr11c chromosomes. Colored blocks indicate locally collinear blocks (LCBs) across the two chromosomes. The intersecting lines between them denote the LCB strikethrough lines.

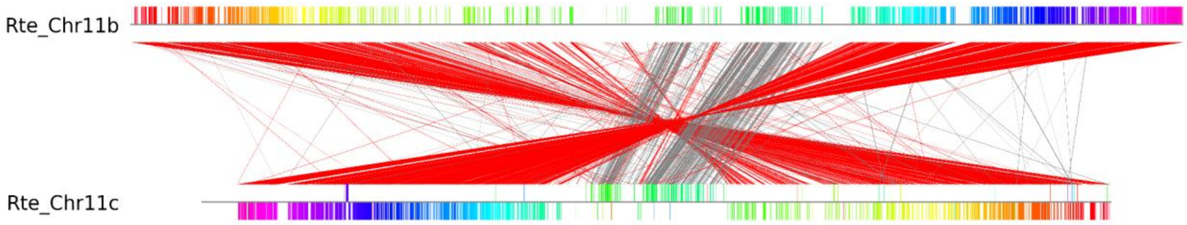


**Supplementary Figure S4: Synteny of 11b and 11c chromosomes in LEZ24 genome:** This figure represents a synteny plot generated by Progressive Mauve between Rte_Chr11b and Rte_Chr11c chromosomes. Colored blocks indicate locally collinear blocks (LCBs) across the two chromosomes. The intersecting lines between them denote the LCB strikethrough lines.

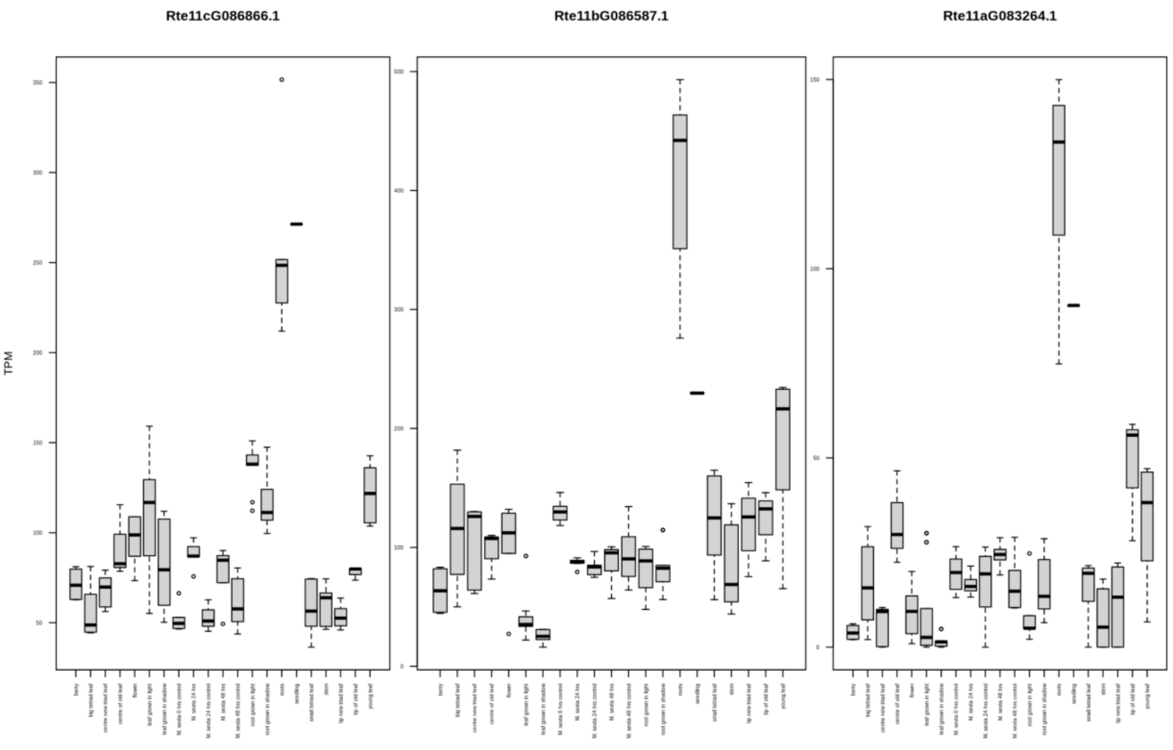


**Supplementary Figure S5: Tissue-specific expression of GS homeologs in LEZ24 genome:** Boxplots representing transcript abundance (TPM) for three GS homeologs- Rte11cG086866.1, Rte11bG086587.1 and Rte11aG083264.1. Distinct expression patterns among the three copies suggest homeolog-specific regulation across different tissue-types, developmental stages and stress treatments (x-axis).

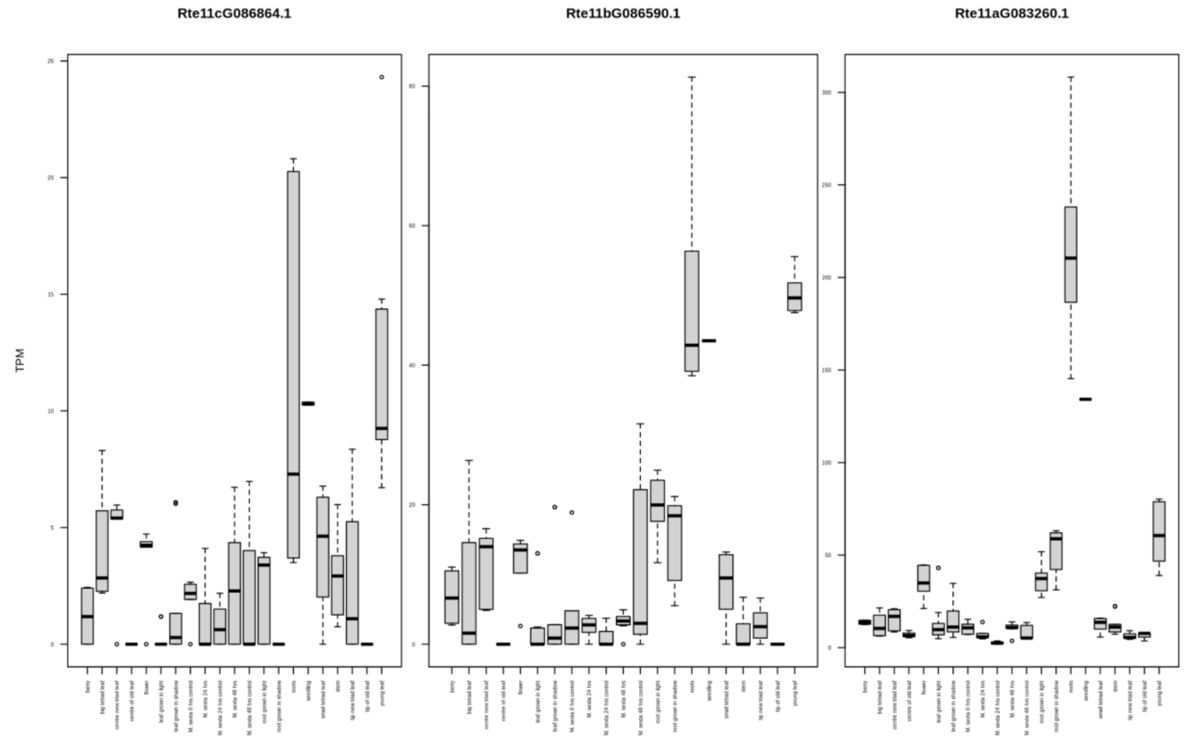


**Supplementary Figure S6: Tissue-specific expression of MSTRG.5530 homeologs in LEZ24 genome:** Boxplots representing transcript abundance (TPM) for three MSTRG.5530 homeologs-Rte11cG086864.1, Rte11bG086590.1 and Rte11aG083260.1. Distinct expression patterns among the three copies suggest homeolog-specific regulation across different tissue-types, developmental stages and stress treatments (x-axis).

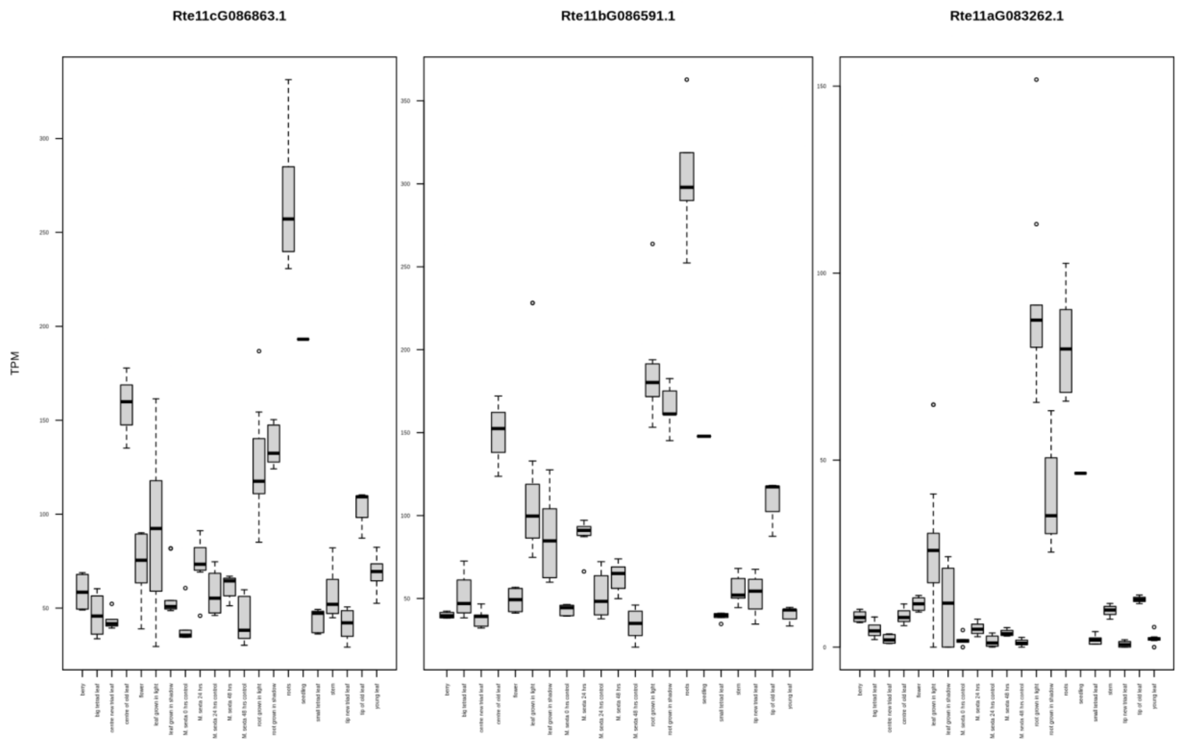


**Supplementary Figure S7: Tissue-specific expression of MSTRG.5531 homeologs in LEZ24 genome:** Boxplots representing transcript abundance (TPM) for three MSTRG.5531 homeologs- Rte11cG086863.1, Rte11bG086591.1 and Rte11aG083262.1. Distinct expression patterns among the three copies suggest homeolog-specific regulation across different tissue-types, developmental stages and stress treatments (x-axis).

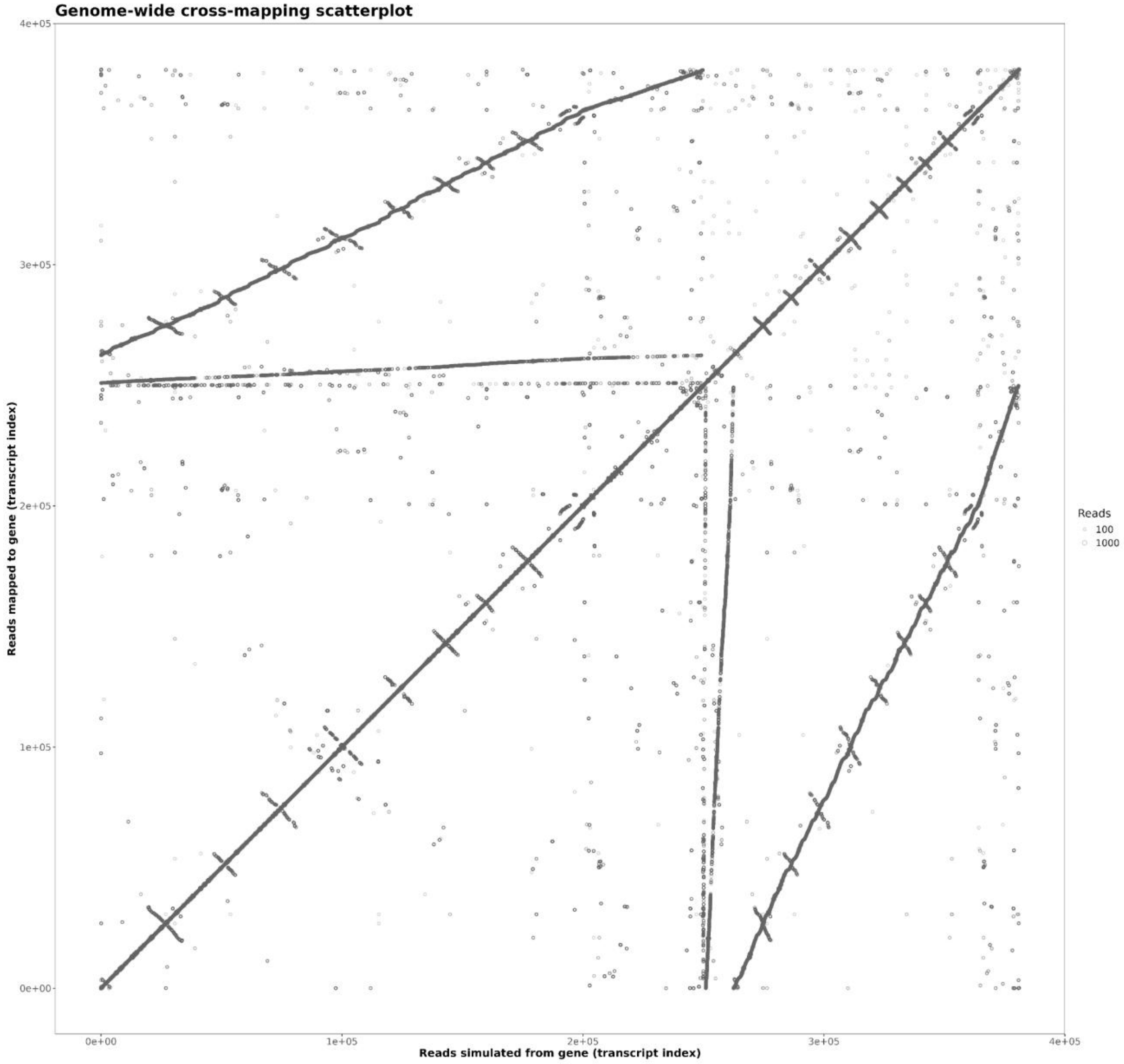


**Supplementary Figure S8: Cross-mapping visualisation for LEZ24 and stringtie transcripts:** Extensive off-diagonal signal among MSTRG models indicating widespread sequence redundancy and high similarity among assembled transcripts.

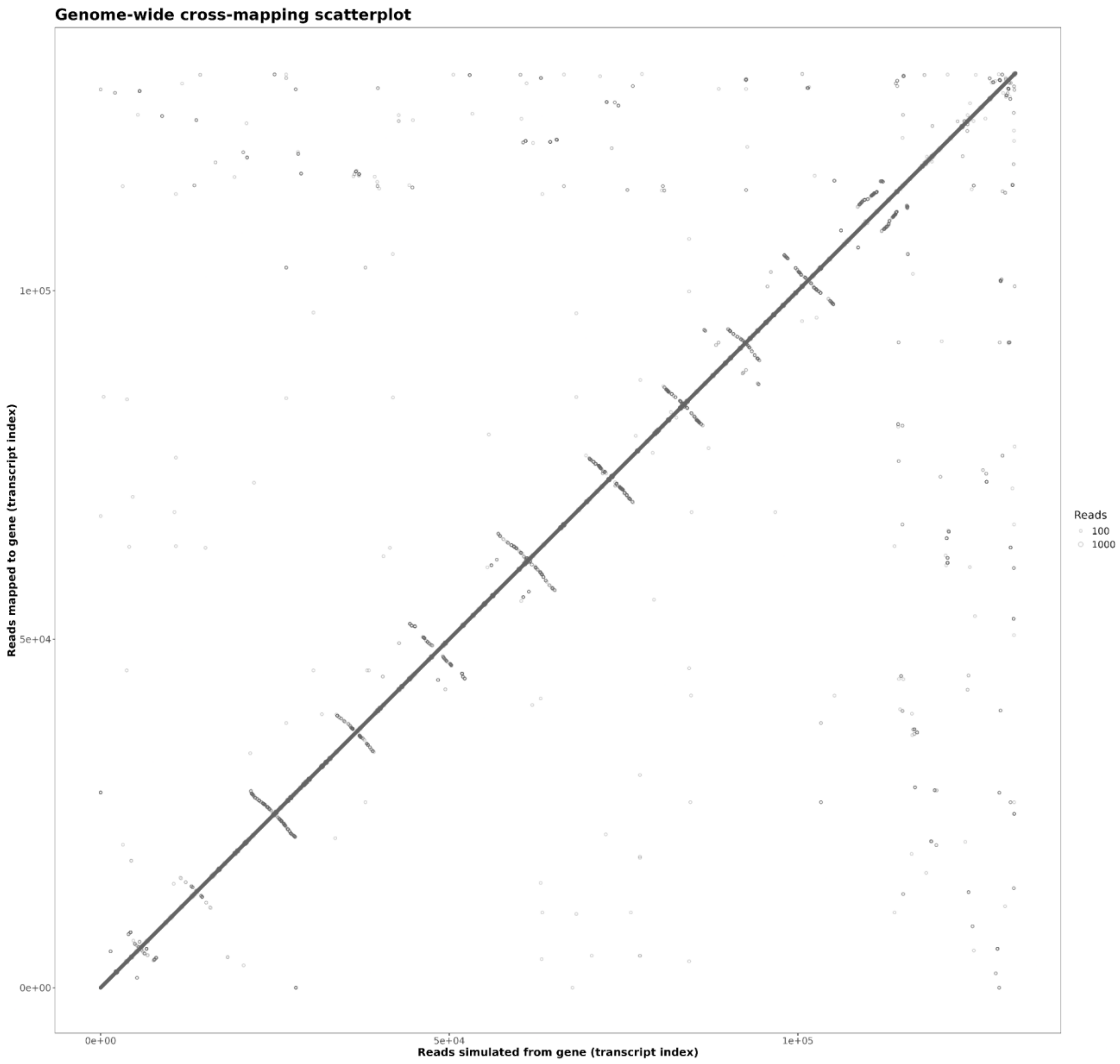


**Supplementary Figure S9: Cross-mapping visualisation for LEZ24 transcripts:** Minimal off-diagonal signal and limited cross-mapping largely confined to isoforms of the same transcript, supporting specificity of read assignment among LEZ24 transcripts.

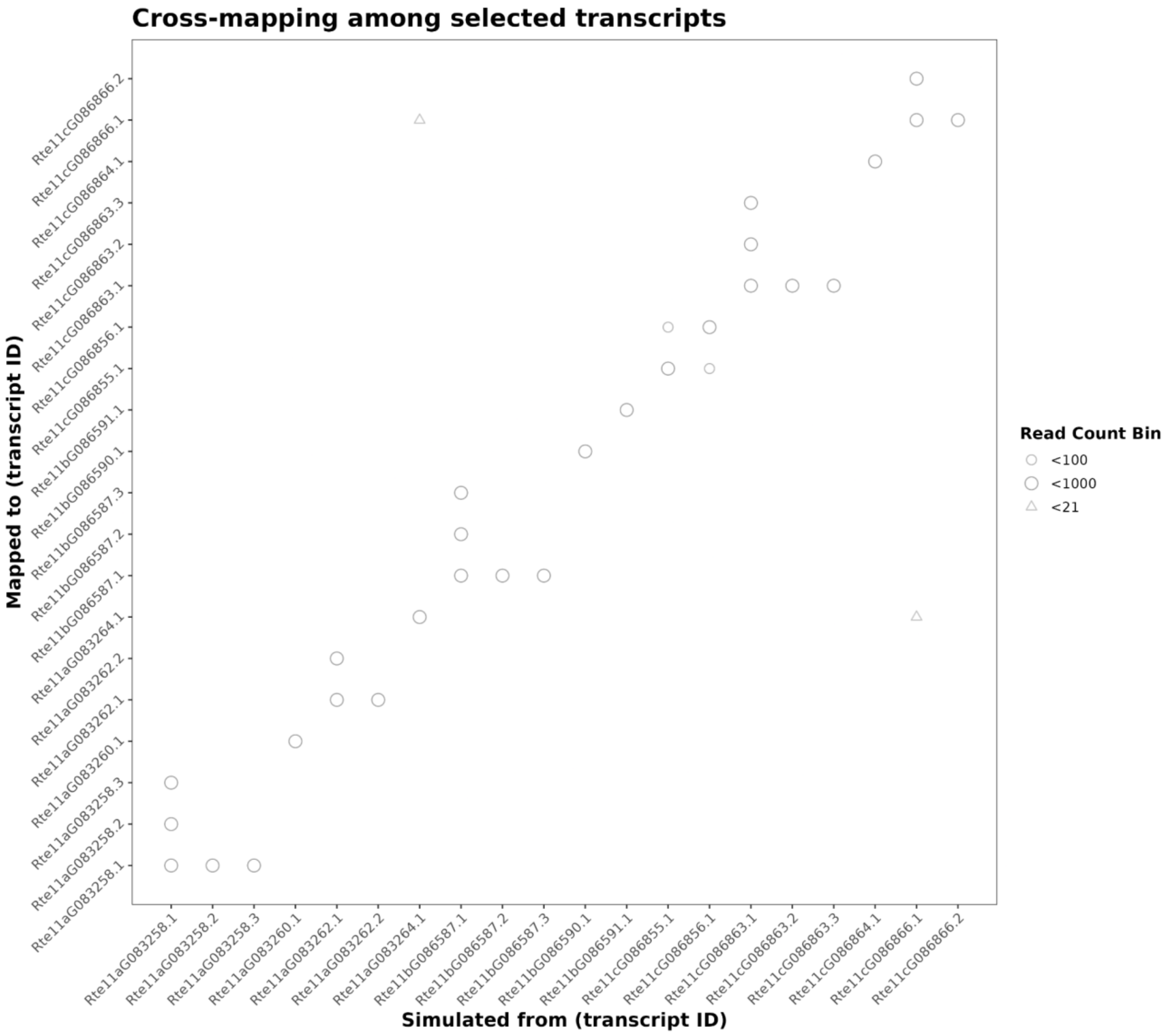


**Supplementary Figure S10: Cross-mapping visualisation for yohimbane candidate transcripts :** Negligible cross-mapping among distinct homeologous copies, supporting the reliability of homeolog-specific expression comparisons in downstream analyses.

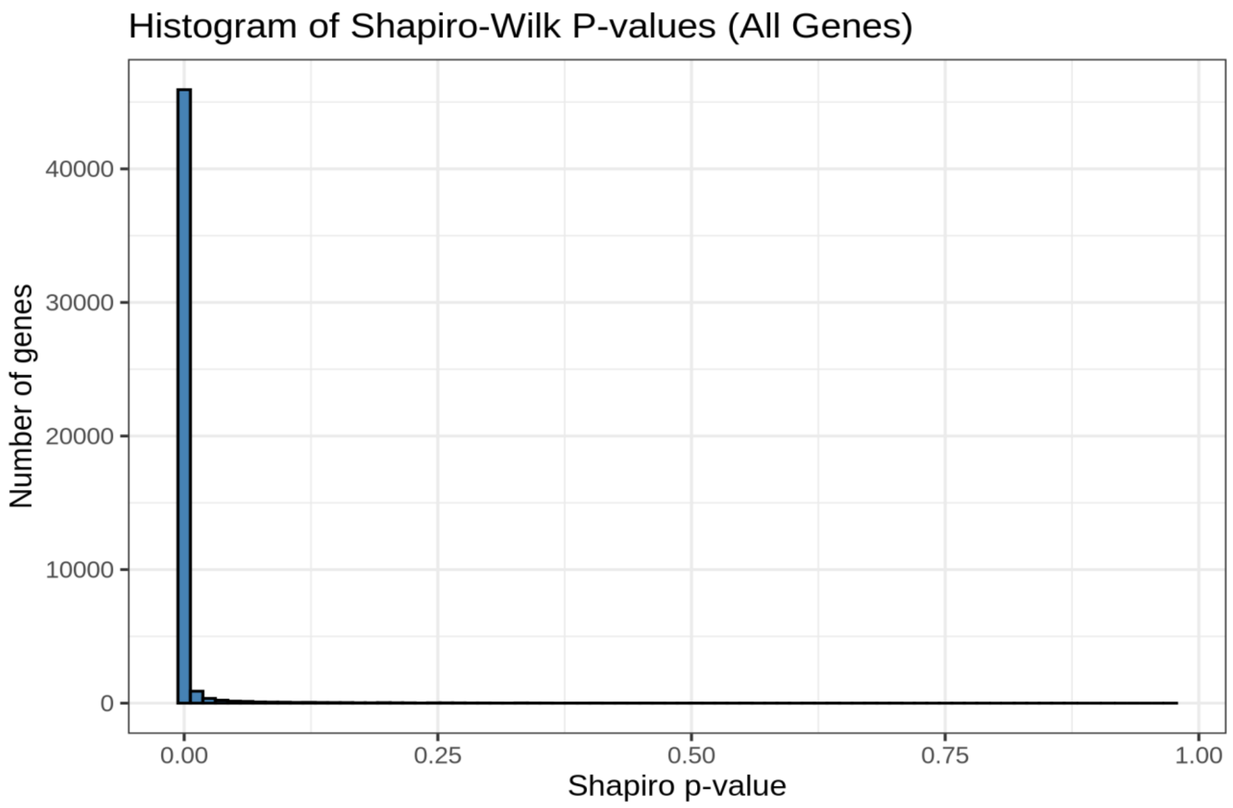


**Supplementary Figure S11: Histogram representing Shapiro–Wilk normality test across genes:** Histogram representing the distribution of Shapiro–Wilk test p-values computed for each gene across the full expression dataset. The x-axis shows Shapiro–Wilk p-values while the y-axis indicates the number of genes falling within each p-value bin. High number of genes with p-values near zero (< 0.05) indicates that expression vectors for the majority of genes deviate significantly from a normal distribution.

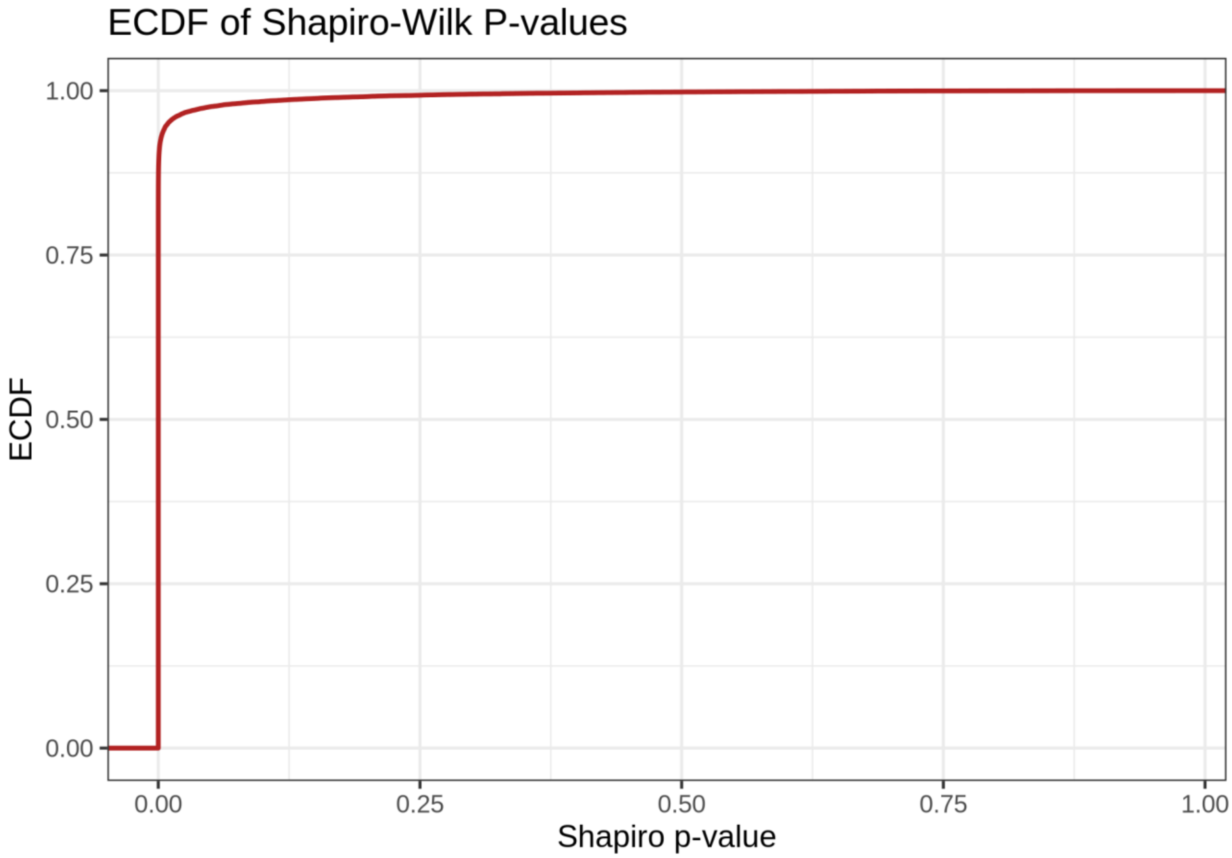


**Supplementary Figure S12: ECDF representing Shapiro–Wilk normality test across genes:** Empirical cumulative distribution function (ECDF) plot representing the cumulative distribution of Shapiro–Wilk test p-values computed for each gene across the full expression dataset. The x-axis shows Shapiro–Wilk p-values, while the y-axis indicates the cumulative fraction of genes with p-values less than or equal to a given threshold. The steep rise near p = 0 shows high percentage of genes with p-values below 0.05 indicating that the expression vectors for majority of genes deviate significantly from a normal distribution.

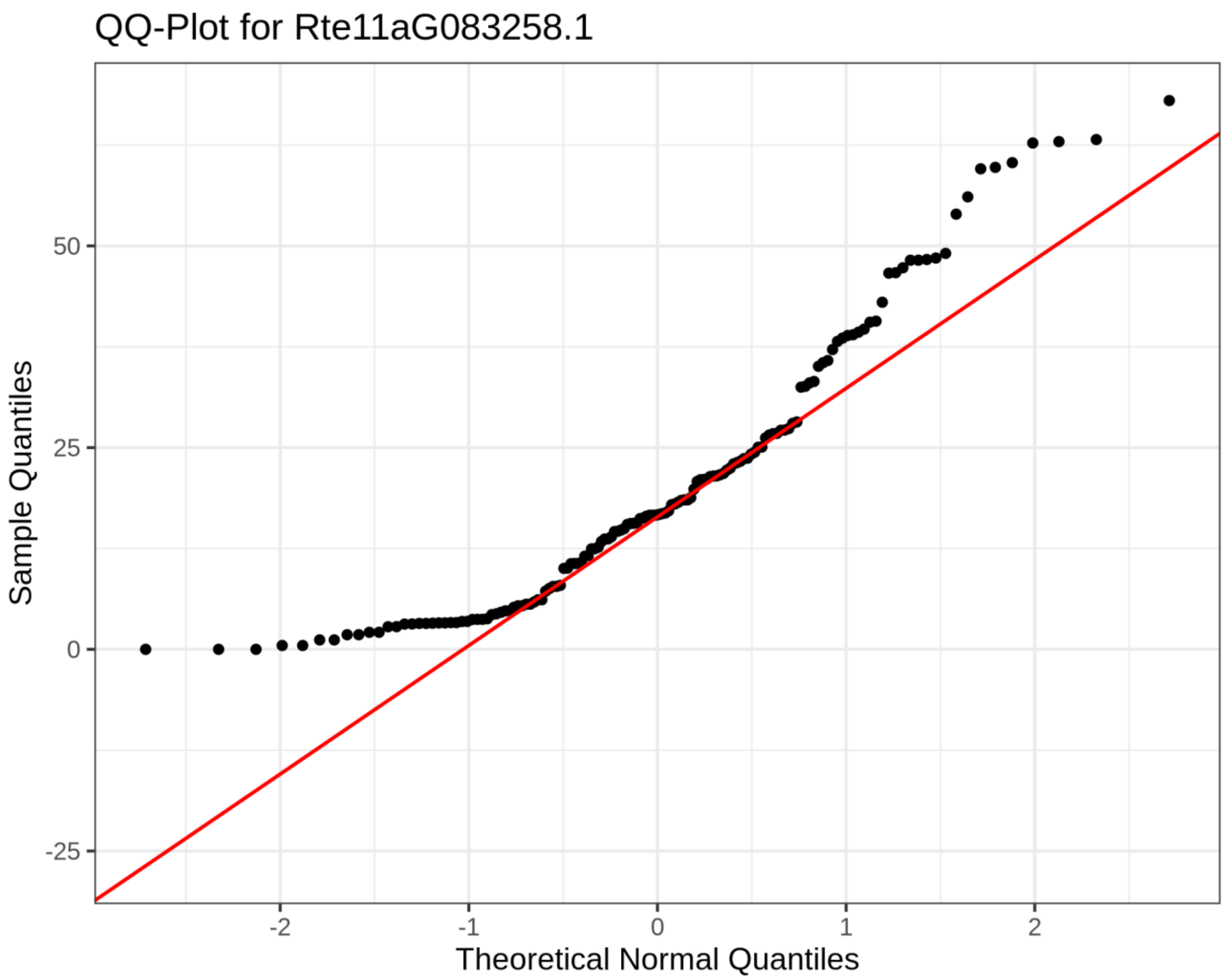


**Supplementary Figure S13: Normality assessment using Quantile–quantile plot for Rte11aG083258.1 transcript:**

Quantile–quantile plot of TPM values for Rte11aG083258.1 across RNA-seq samples in the transcription expression vector compared against the expected quantiles of a normal distribution. The red diagonal line represents the theoretical relationship under normality. Deviation of observed points from the line, particularly the low quantiles near zero and upward curvature at the upper tail, indicates that the transcript expression values are not normally distributed across tissues.

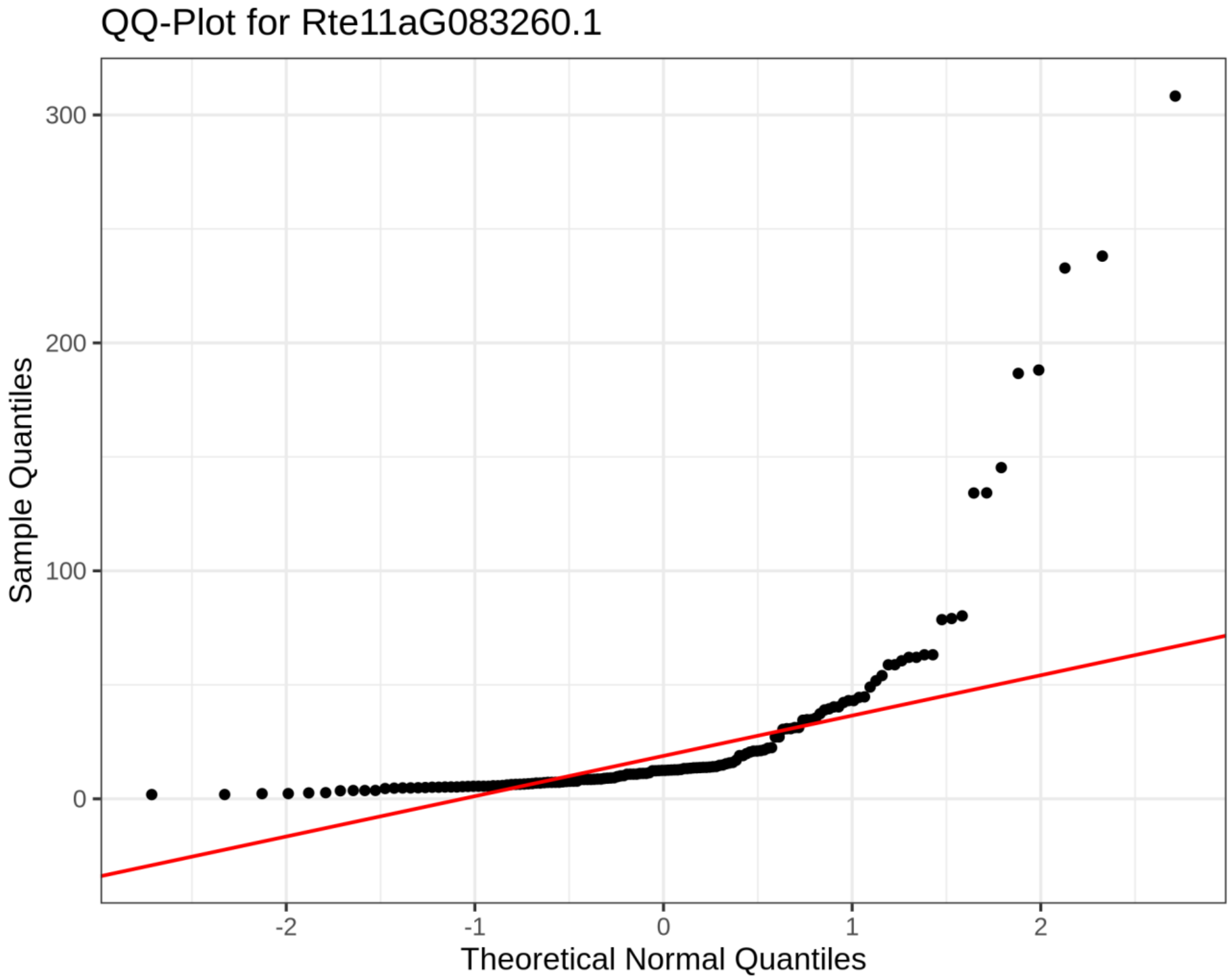


**Supplementary Figure S14: Normality assessment using Quantile–quantile plot for Rte11aG083260.1 transcript:**

Quantile–quantile plot of TPM values for Rte11aG083260.1 across RNA-seq samples in the transcription expression vector compared against the expected quantiles of a normal distribution. The red diagonal line represents the theoretical relationship under normality. Deviation of observed points from the line, particularly the low quantiles near zero and upward curvature at the upper tail, indicates that the transcript expression values are not normally distributed across tissues.

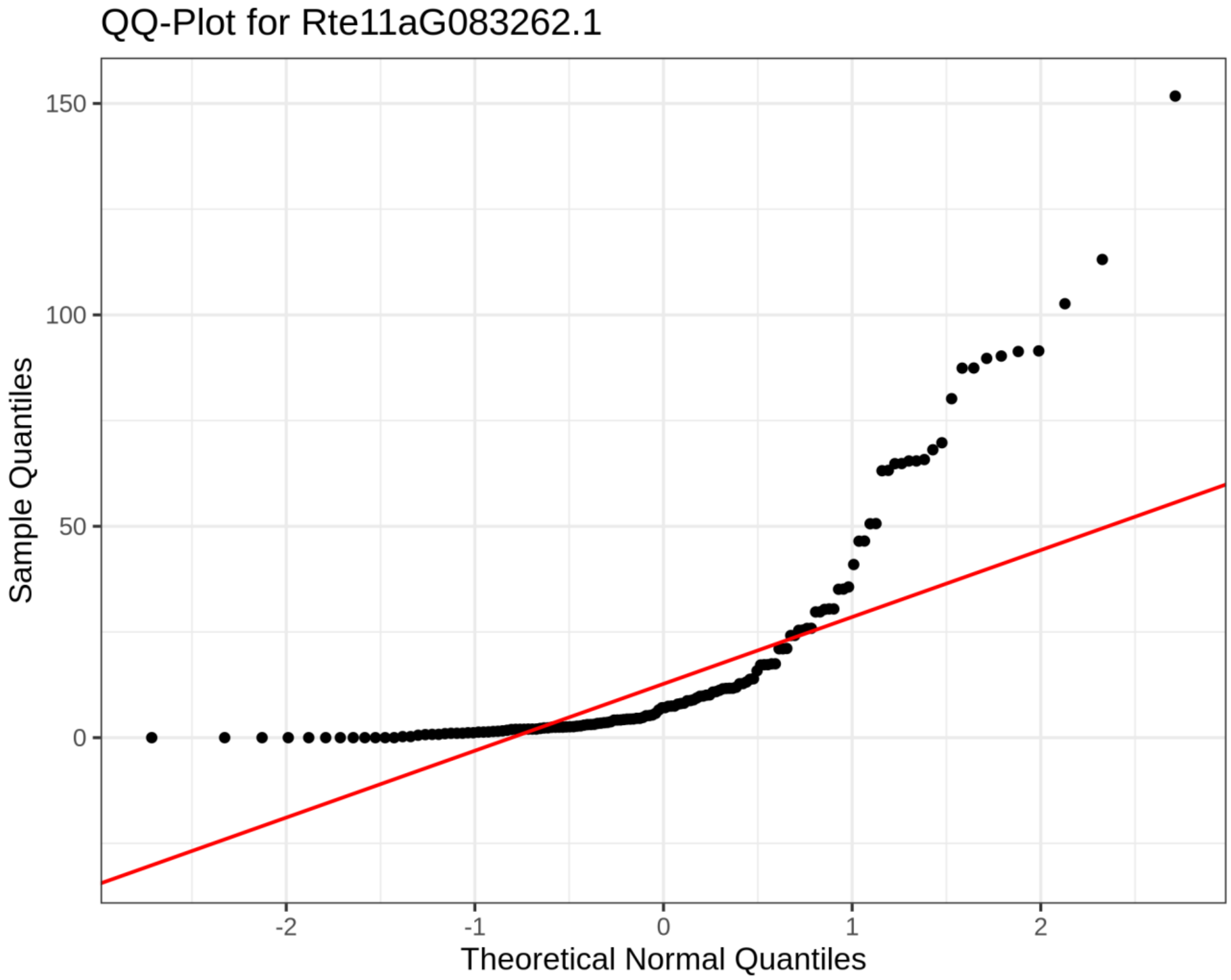


**Supplementary Figure S15: Normality assessment using Quantile–quantile plot for Rte11aG083262.1 transcript:**

Quantile–quantile plot of TPM values for Rte11aG083262.1 across RNA-seq samples in the transcription expression vector compared against the expected quantiles of a normal distribution. The red diagonal line represents the theoretical relationship under normality. Deviation of observed points from the line, particularly the low quantiles near zero and upward curvature at the upper tail, indicates that the transcript expression values are not normally distributed across tissues.

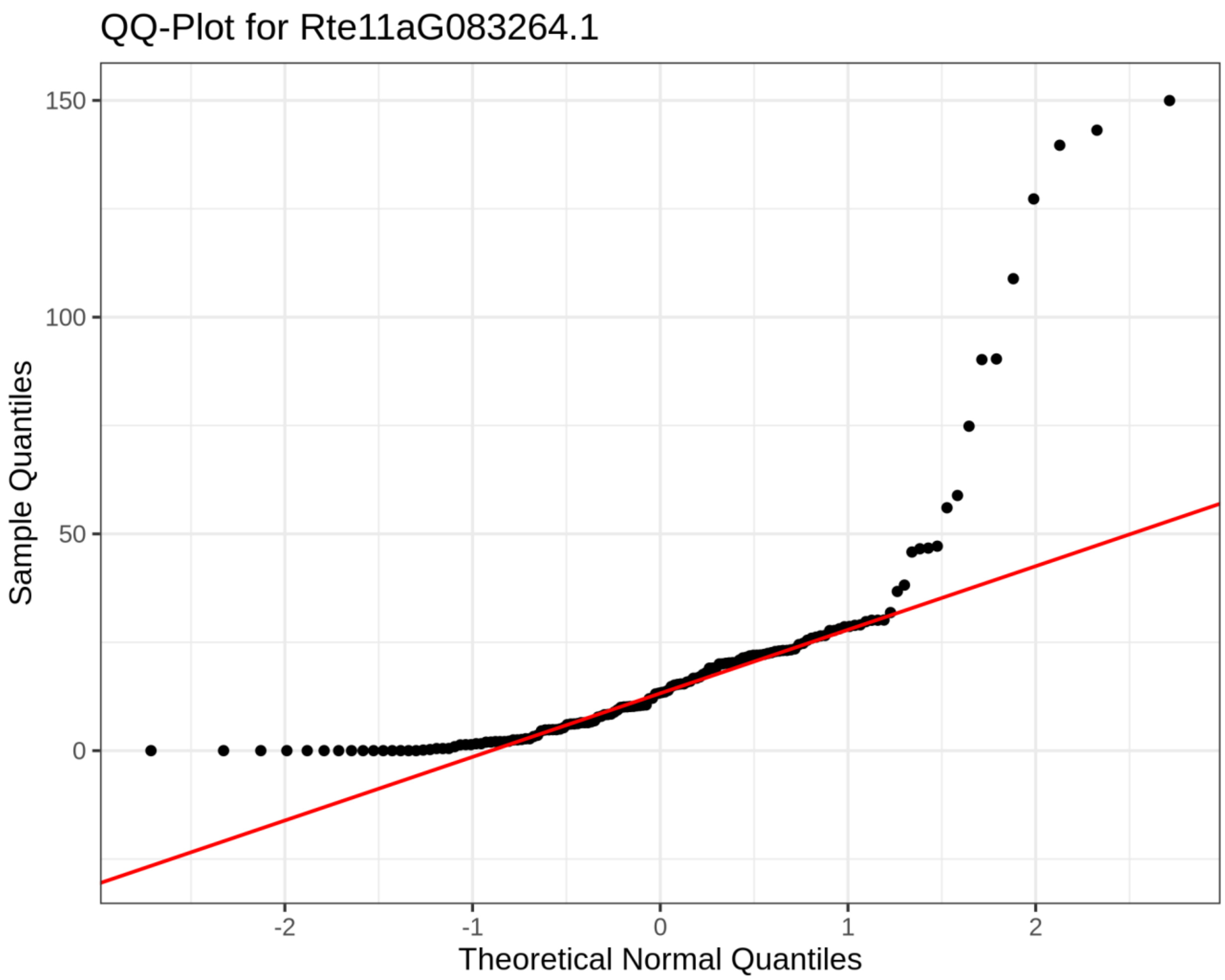


**Supplementary Figure S16: Normality assessment using Quantile–quantile plot for Rte11aG083264.1 transcript:**

Quantile–quantile plot of TPM values for Rte11aG083264.1 across RNA-seq samples in the transcription expression vector compared against the expected quantiles of a normal distribution. The red diagonal line represents the theoretical relationship under normality. Deviation of observed points from the line, particularly the low quantiles near zero and upward curvature at the upper tail, indicates that the transcript expression values are not normally distributed across tissues.

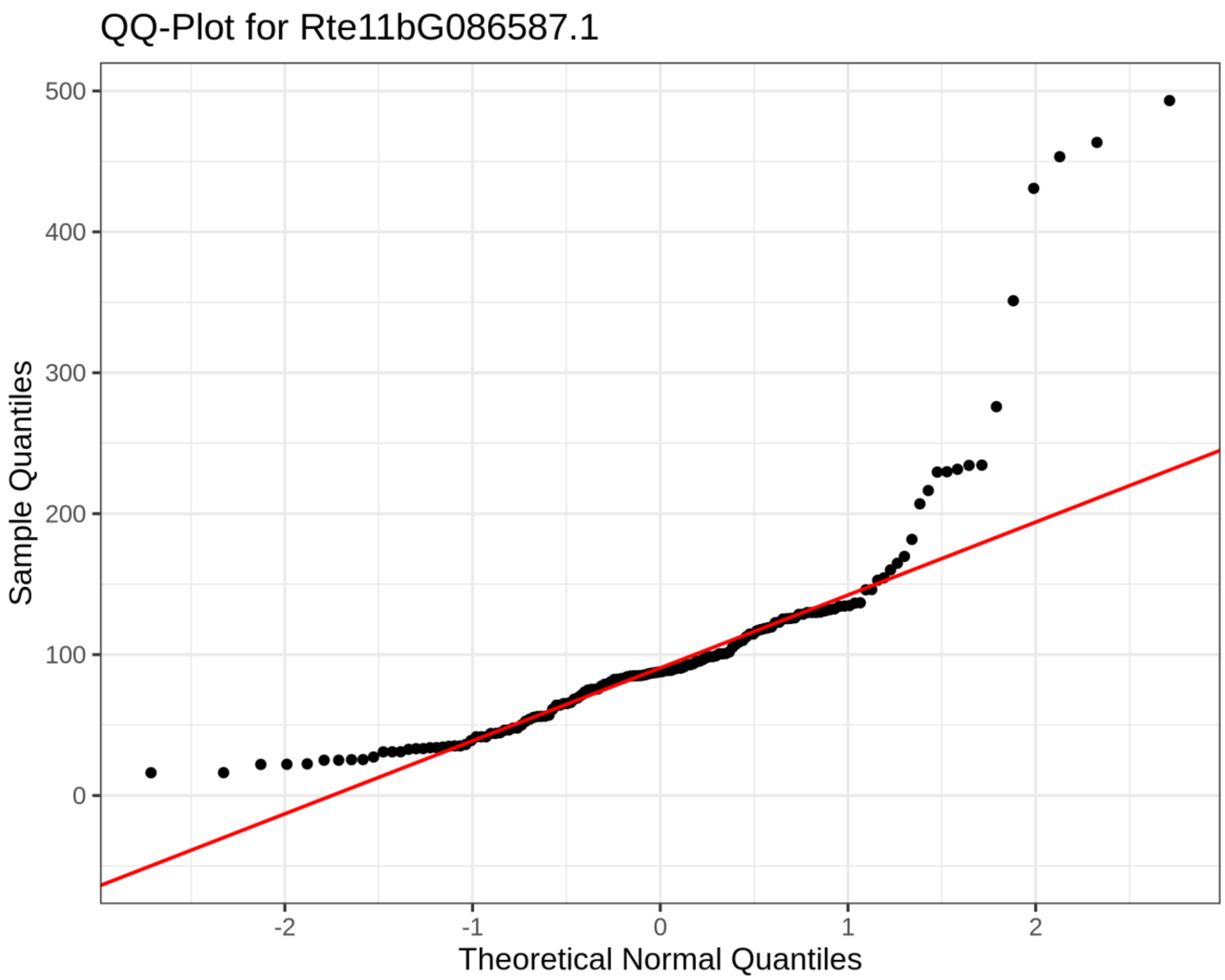


**Supplementary Figure S17: Normality assessment using Quantile–quantile plot for Rte11bG086587.1 transcript:**

Quantile–quantile plot of TPM values for Rte11bG086587.1 across RNA-seq samples in the transcription expression vector compared against the expected quantiles of a normal distribution. The red diagonal line represents the theoretical relationship under normality. Deviation of observed points from the line, particularly the low quantiles near zero and upward curvature at the upper tail, indicates that the transcript expression values are not normally distributed across tissues.

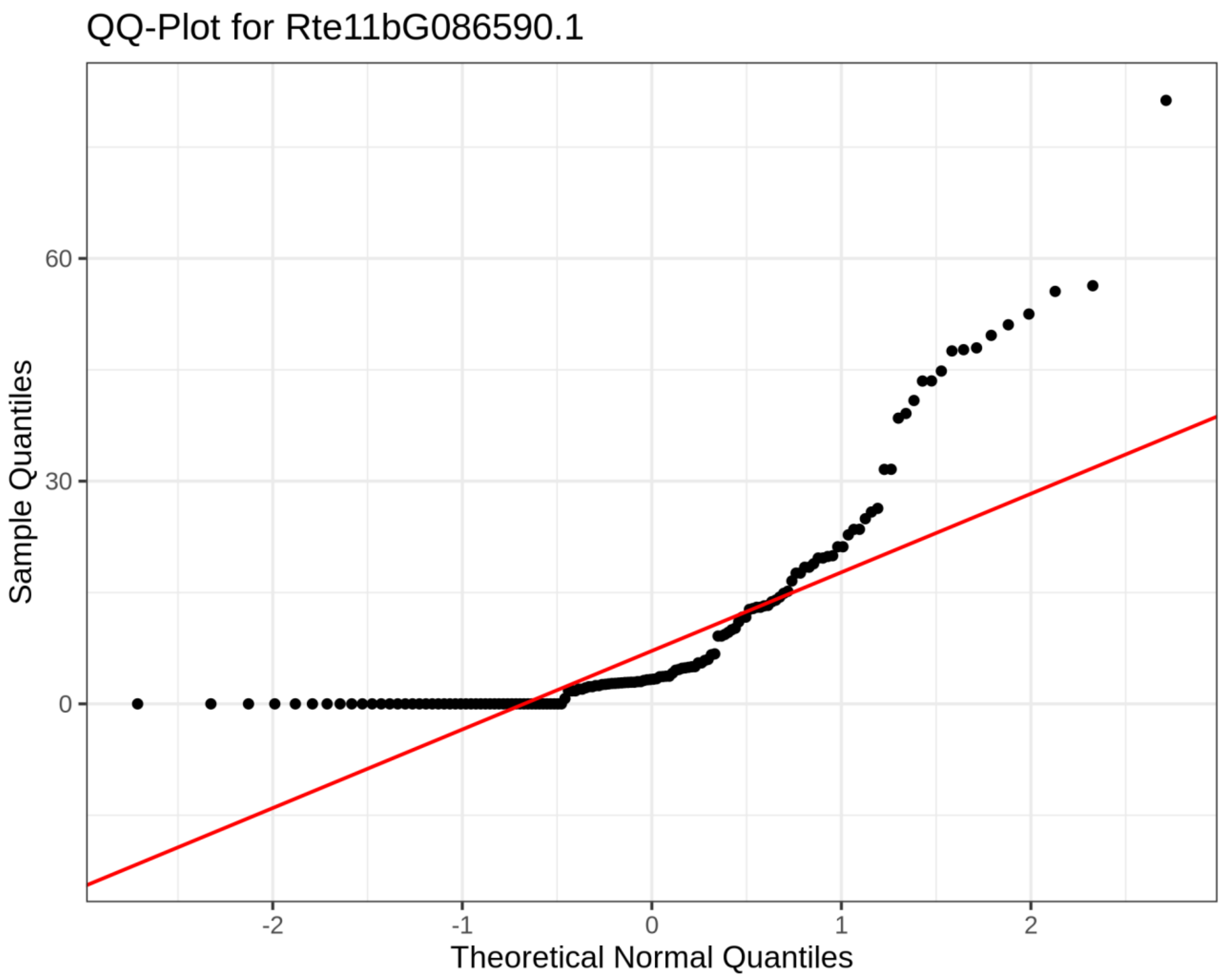


**Supplementary Figure S18: Normality assessment using Quantile–quantile plot for Rte11bG086590.1 transcript:**

Quantile–quantile plot of TPM values for Rte11bG086590.1 across RNA-seq samples in the transcription expression vector compared against the expected quantiles of a normal distribution. The red diagonal line represents the theoretical relationship under normality. Deviation of observed points from the line, particularly the low quantiles near zero and upward curvature at the upper tail, indicates that the transcript expression values are not normally distributed across tissues.

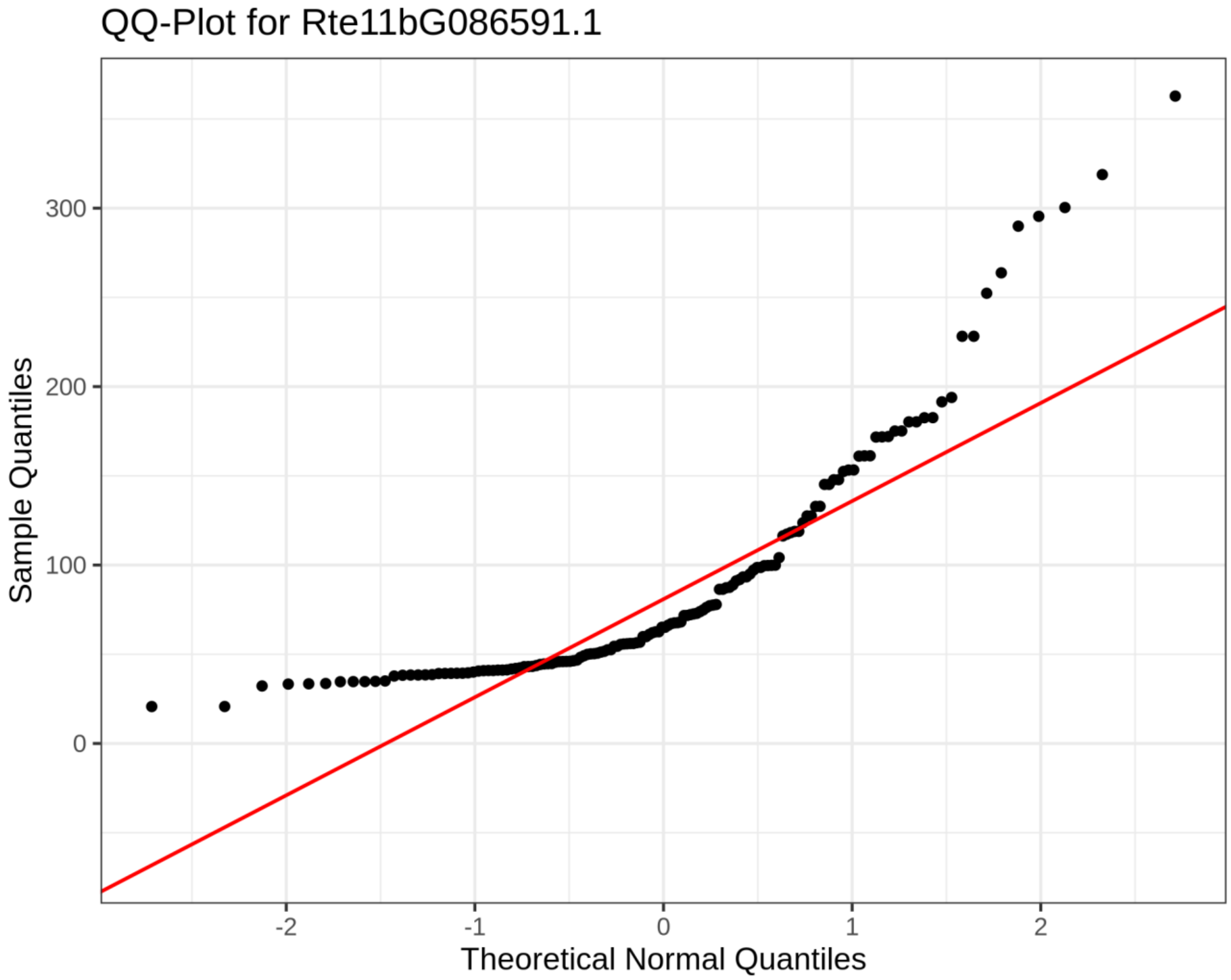


**Supplementary Figure S19: Normality assessment using Quantile–quantile plot for Rte11bG086591.1 transcript:**

Quantile–quantile plot of TPM values for Rte11bG086591.1 across RNA-seq samples in the transcription expression vector compared against the expected quantiles of a normal distribution. The red diagonal line represents the theoretical relationship under normality. Deviation of observed points from the line, particularly the low quantiles near zero and upward curvature at the upper tail, indicates that the transcript expression values are not normally distributed across tissues.

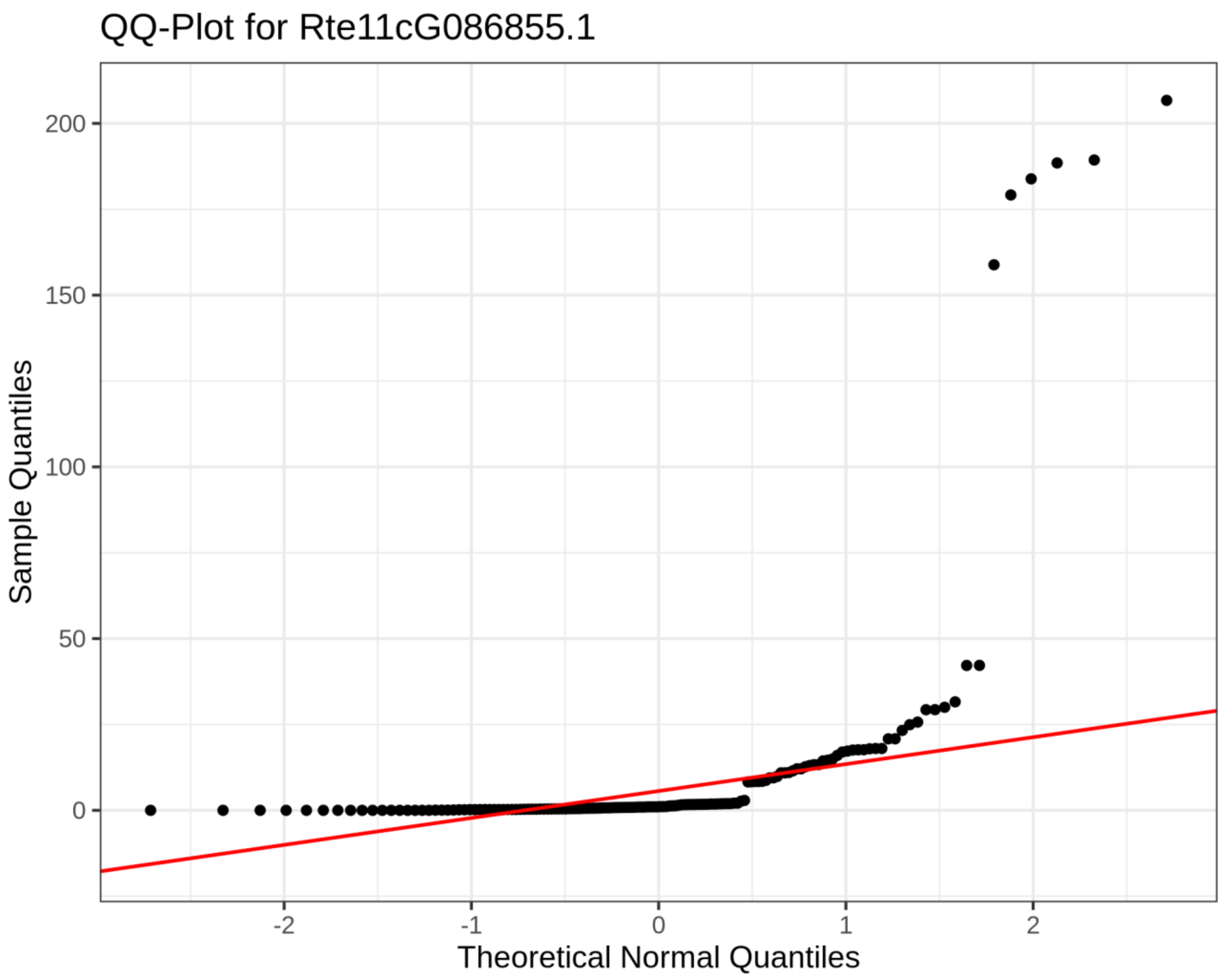


**Supplementary Figure S20: Normality assessment using Quantile–quantile plot for Rte11cG086855.1 transcript:**

Quantile–quantile plot of TPM values for Rte11cG086855.1 across RNA-seq samples in the transcription expression vector compared against the expected quantiles of a normal distribution. The red diagonal line represents the theoretical relationship under normality. Deviation of observed points from the line, particularly the low quantiles near zero and upward curvature at the upper tail, indicates that the transcript expression values are not normally distributed across tissues.

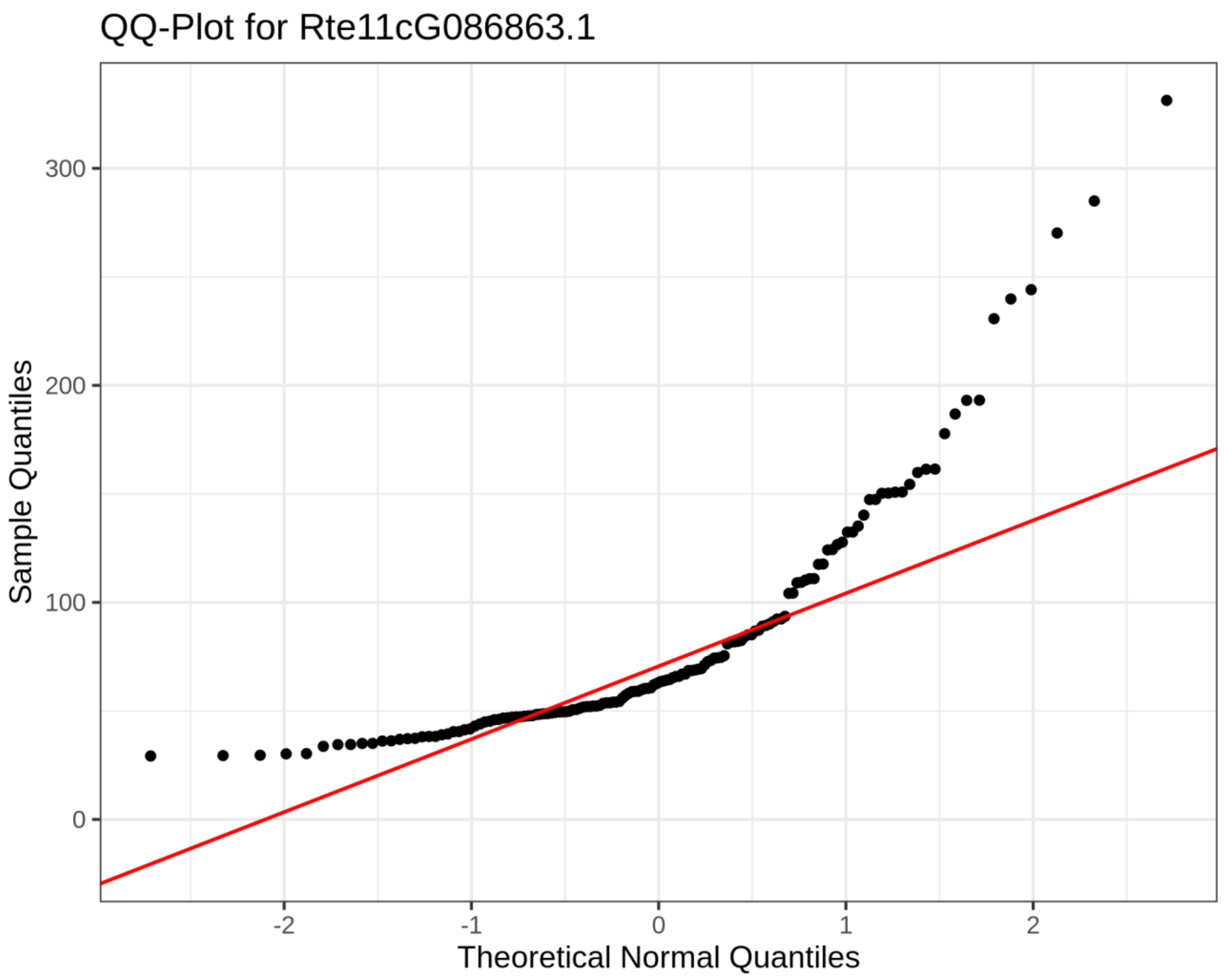


**Supplementary Figure S21: Normality assessment using Quantile–quantile plot for Rte11cG086863.1 transcript:**

Quantile–quantile plot of TPM values for Rte11cG086863.1 across RNA-seq samples in the transcription expression vector compared against the expected quantiles of a normal distribution. The red diagonal line represents the theoretical relationship under normality. Deviation of observed points from the line, particularly the low quantiles near zero and upward curvature at the upper tail, indicates that the transcript expression values are not normally distributed across tissues.

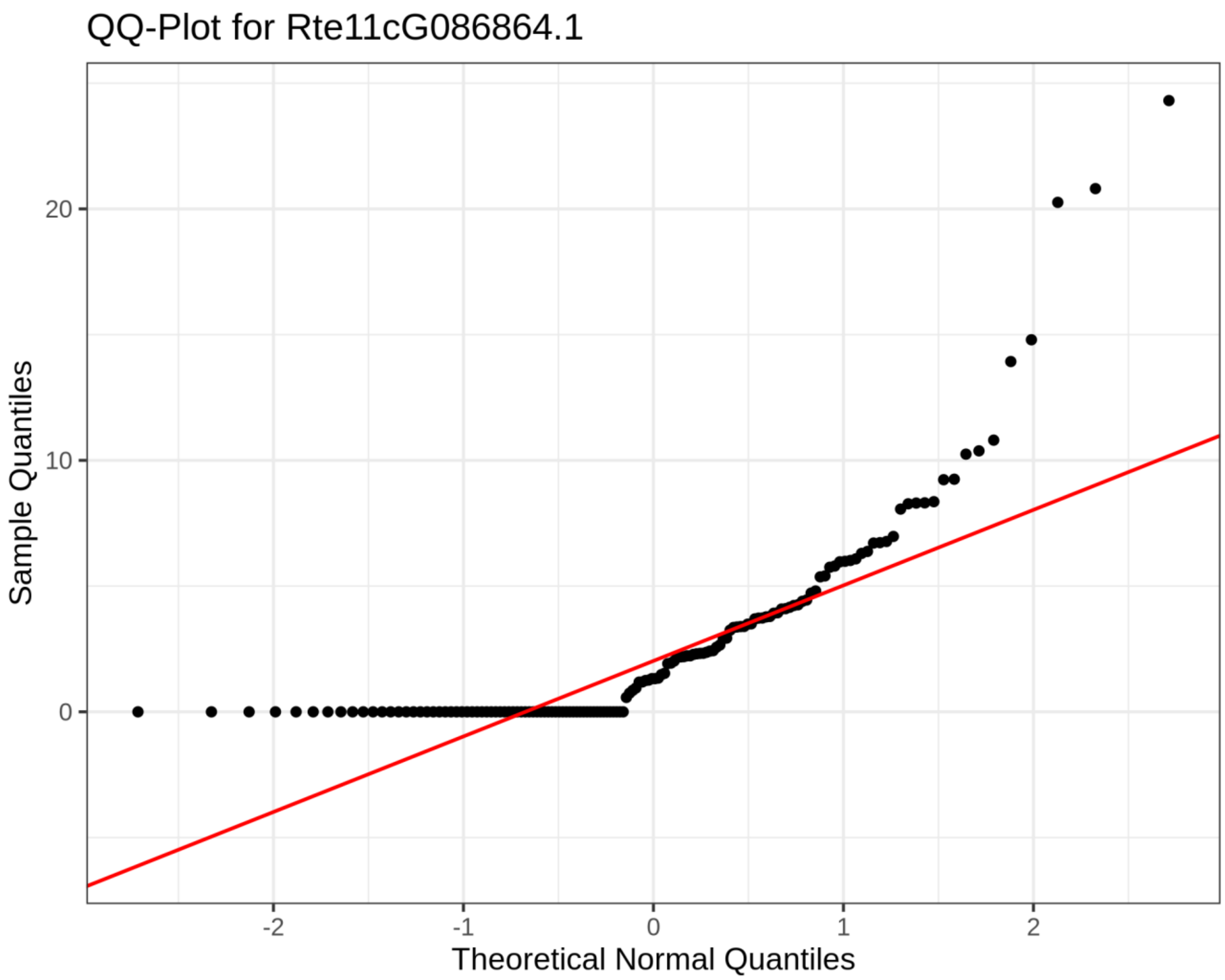


**Supplementary Figure S22: Normality assessment using Quantile–quantile plot for Rte11cG086864.1 transcript:**

Quantile–quantile plot of TPM values for Rte11cG086864.1 across RNA-seq samples in the transcription expression vector compared against the expected quantiles of a normal distribution. The red diagonal line represents the theoretical relationship under normality. Deviation of observed points from the line, particularly the low quantiles near zero and upward curvature at the upper tail, indicates that the transcript expression values are not normally distributed across tissues.

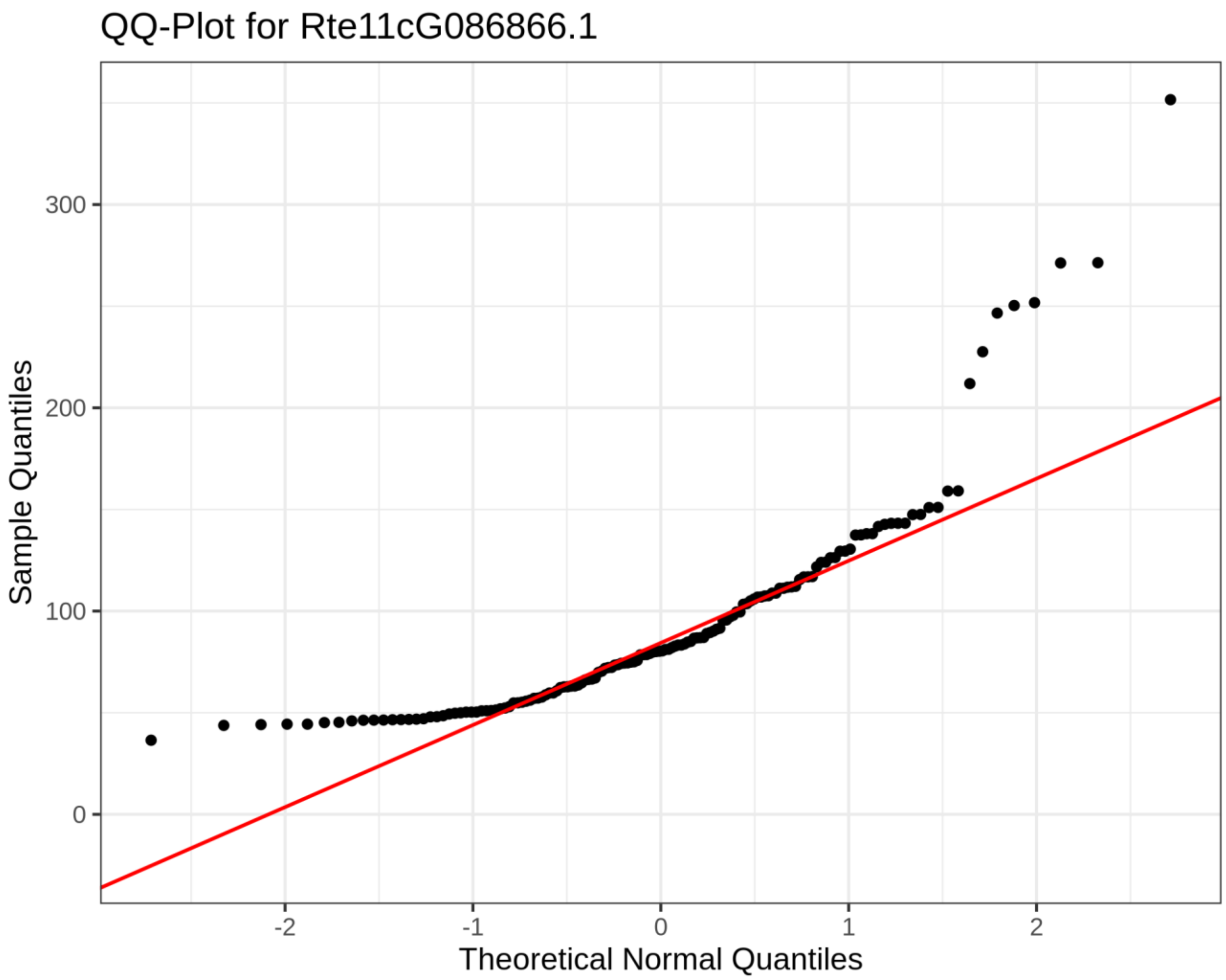


**Supplementary Figure S23: Normality assessment using Quantile–quantile plot for Rte11cG086866.1 transcript:**

Quantile–quantile plot of TPM values for Rte11cG086866.1 across RNA-seq samples in the transcription expression vector compared against the expected quantiles of a normal distribution. The red diagonal line represents the theoretical relationship under normality. Deviation of observed points from the line, particularly the low quantiles near zero and upward curvature at the upper tail, indicates that the transcript expression values are not normally distributed across tissues.

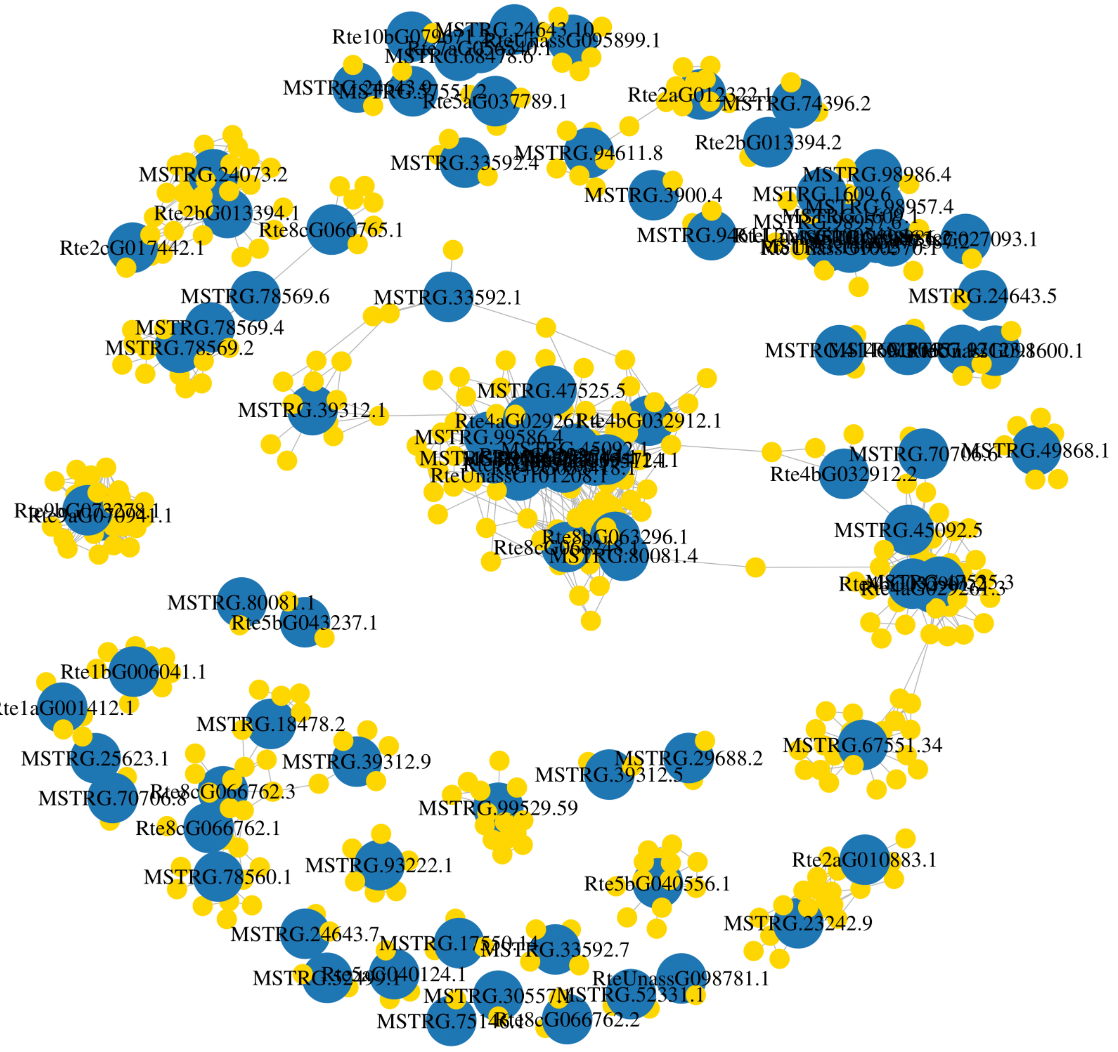


**Supplementary Figure S24: MIA co-expression network:** Transcripts are represented by nodes, connected by edges when their HRR value was below 30.

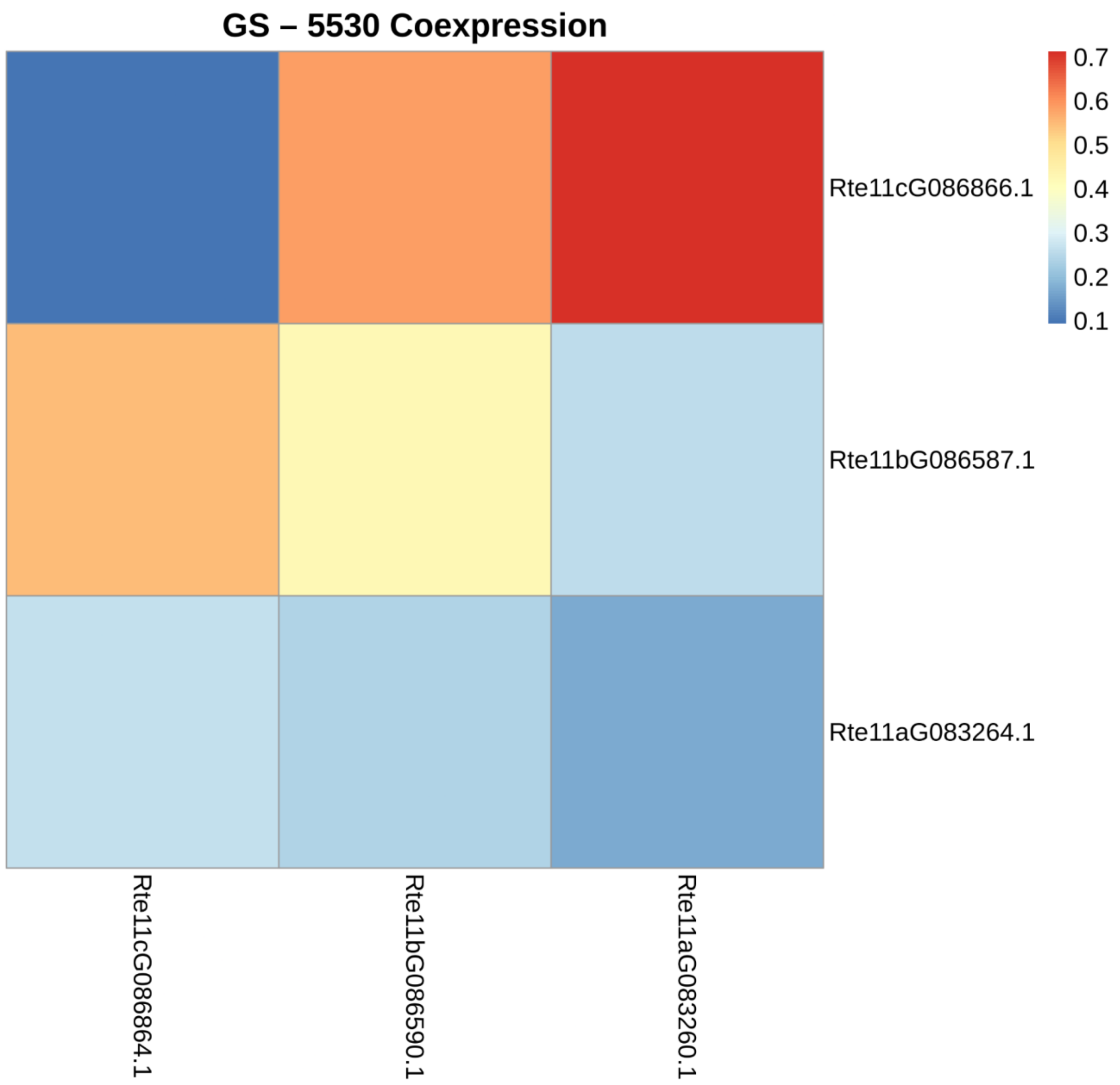


**Supplementary Figure S25: Co-expression between GS and MSTRG.5530 WGT-derived copies in the LEZ24 genome:** Heatmap representing coexpression patterns among homeologs of GS (x-axis) and MSTRG.5530 (y-axis) in the LEZ24 genome. Each square-block represents the Spearman's rank correlation coefficient of expression profiles across the RNA-seq expression atlas between the corresponding transcript pairs. Correlation strength is indicated by colour intensity as shown by the scale bar on the right.

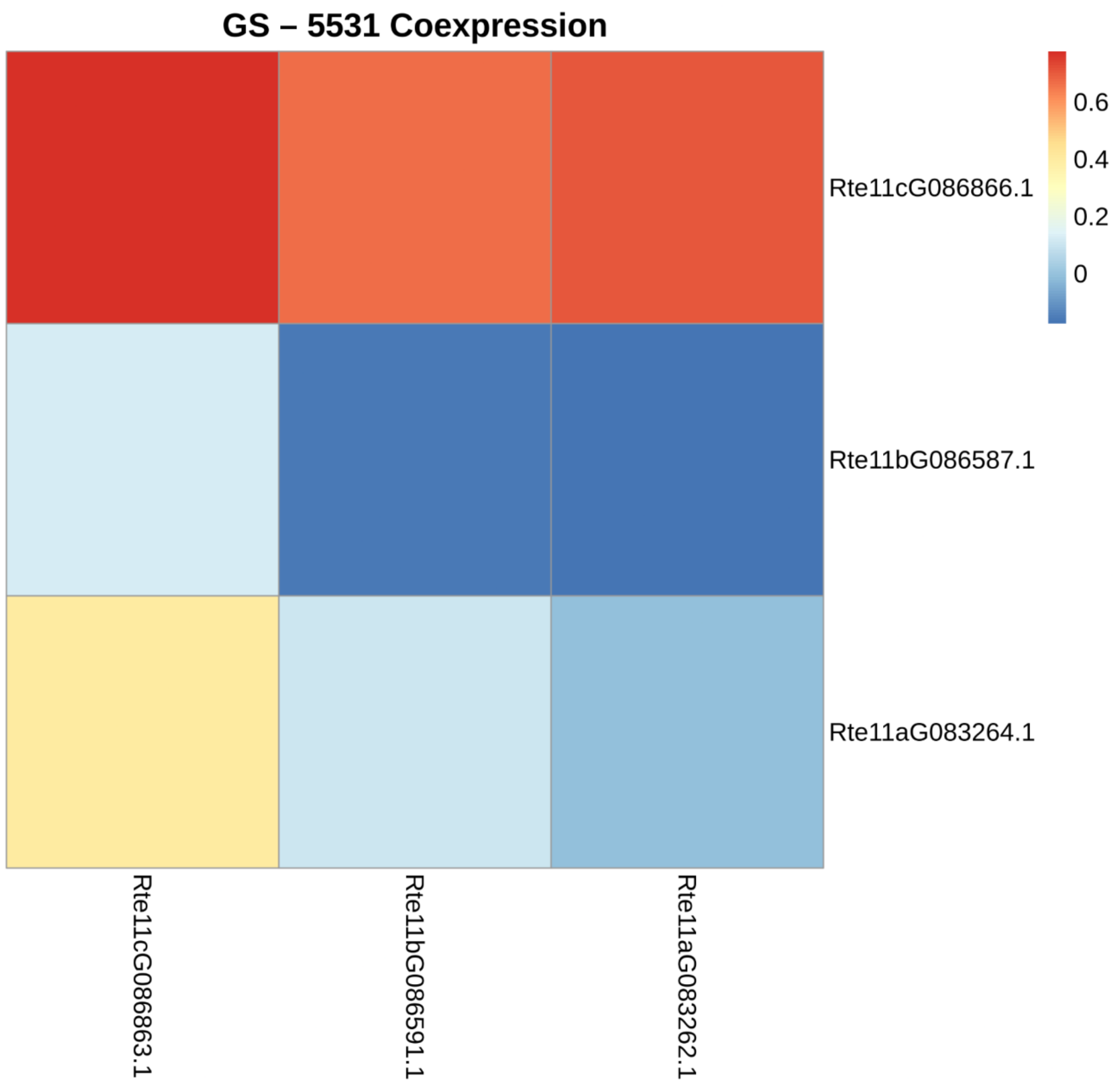


**Supplementary Figure S26: Co-expression between GS and MSTRG.5531 WGT-derived copies in the LEZ24 genome:** Heatmap representing coexpression patterns among homeologs of GS (x-axis) and MSTRG.5531 (y-axis) in the LEZ24 genome. Each square-block represents the Spearman's rank correlation coefficient of expression profiles across the RNA-seq expression atlas between the corresponding transcript pairs. Correlation strength is indicated by colour intensity as shown by the scale bar on the right.

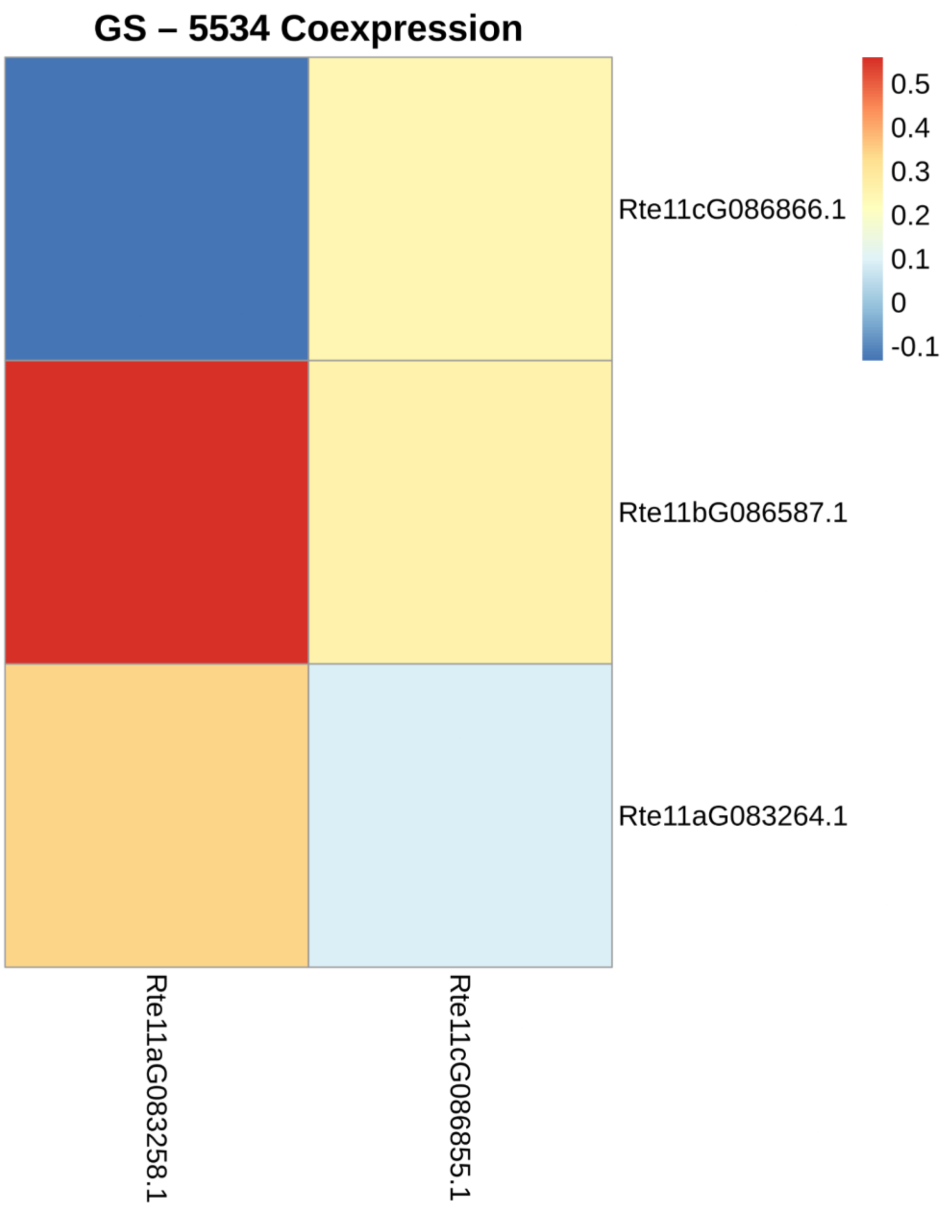


**Supplementary Figure S27: Co-expression between GS and MSTRG.5534 WGT-derived copies in the LEZ24 genome:** Heatmap representing coexpression patterns among homeologs of GS (x-axis) and MSTRG.55304 (y-axis) in the LEZ24 genome. Each square-block represents the Spearman's rank correlation coefficients of expression profiles across the RNA-seq expression atlas between the corresponding transcript paris. Correlation strength is indicated by colour intensity as shown by the scale bar on the right.

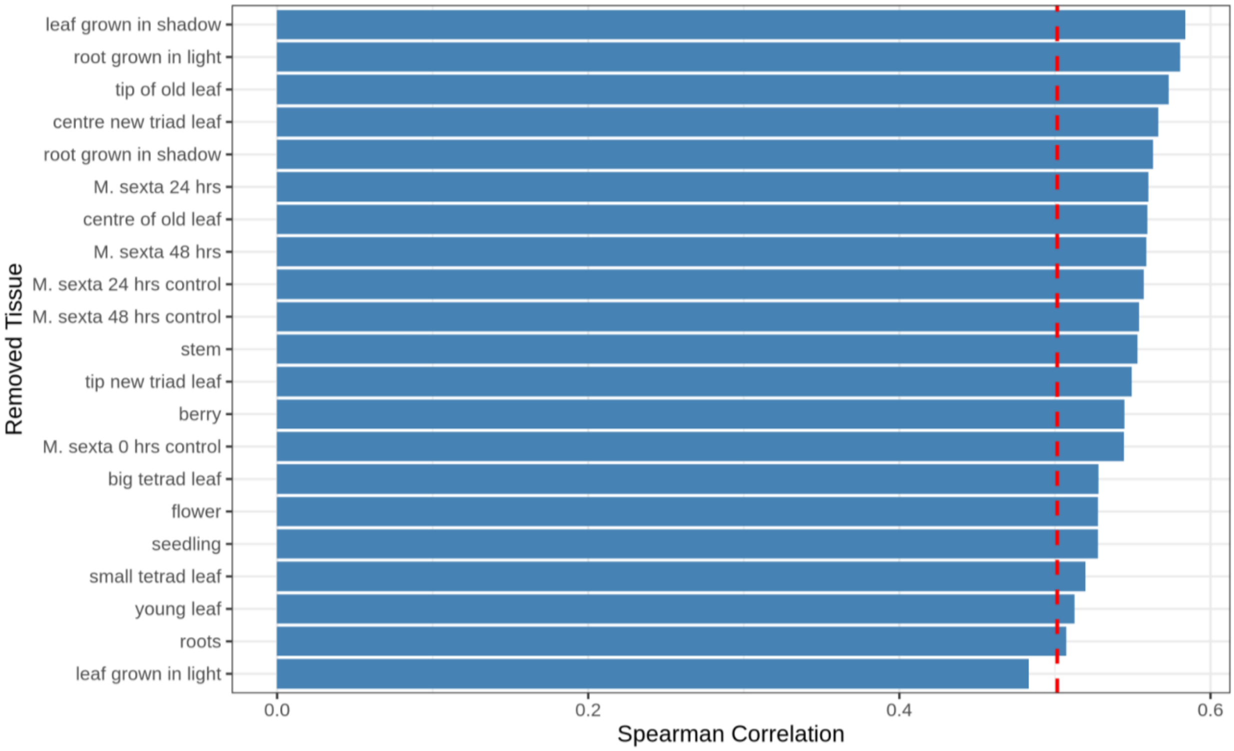


**Supplementary Figure S28: Leave-one-tissue-out (LOTO) correlation stability analysis between GS-11b and MSTRG.5531-11c homeolog across tissues:** Horizontal bar plot showing Spearman correlation coefficients between the GS-11b (Rte11bG086587.1) and MSTRG.5530-11c (Rte11cG086864.1) homeolog copies after iteratively excluding all samples from one tissue type at a time.

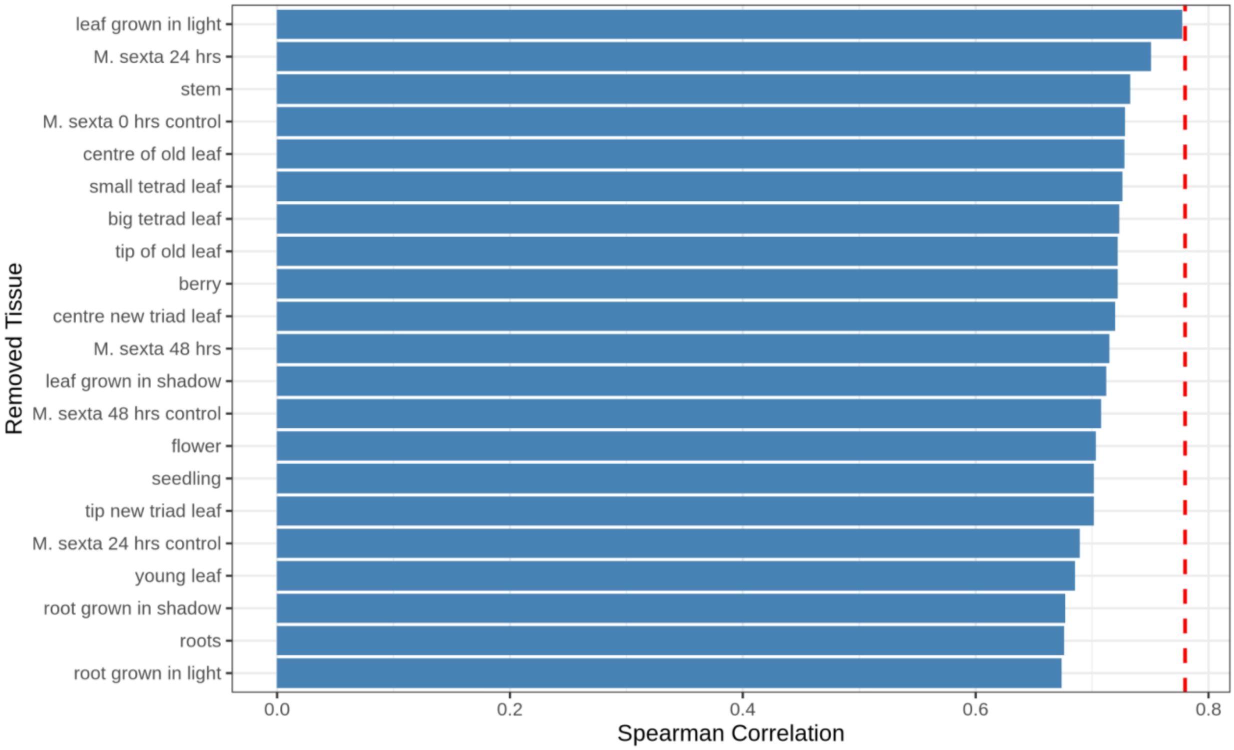


**Supplementary Figure S29: Leave-one-tissue-out (LOTO) correlation stability analysis between GS-11c and MSTRG.5530-11a homeolog across tissues:** Horizontal bar plot showing Spearman correlation coefficients between the GS-11c (Rte11cG086866.1) and MSTRG.5530-11a (Rte11aG083260.1) homeolog copies after iteratively excluding all samples from one tissue type at a time.

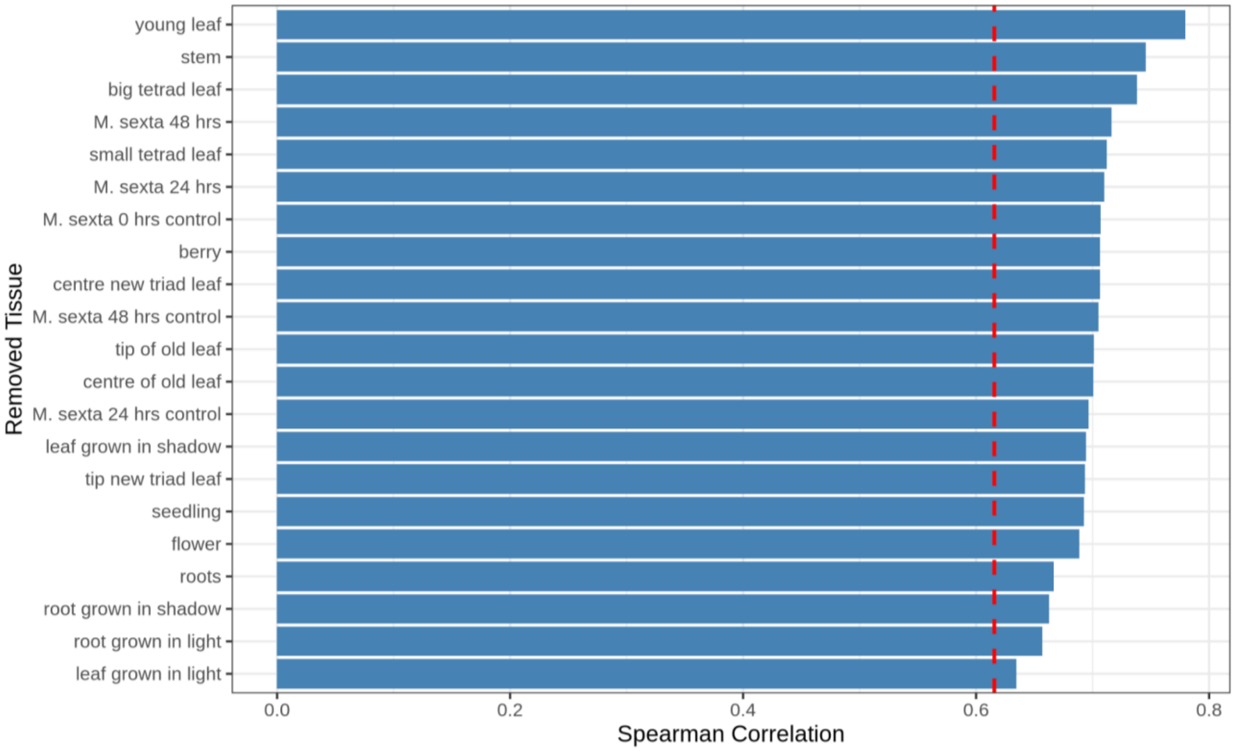


**Supplementary Figure S30: Leave-one-tissue-out (LOTO) correlation stability analysis between GS-11c and MSTRG.5531-11a homeolog across tissues:** Horizontal bar plot showing Spearman correlation coefficients between the GS-11c (Rte11cG086866.1) and MSTRG.5531-11a (Rte11aG083262.1) homeolog copies after iteratively excluding all samples from one tissue type at a time.

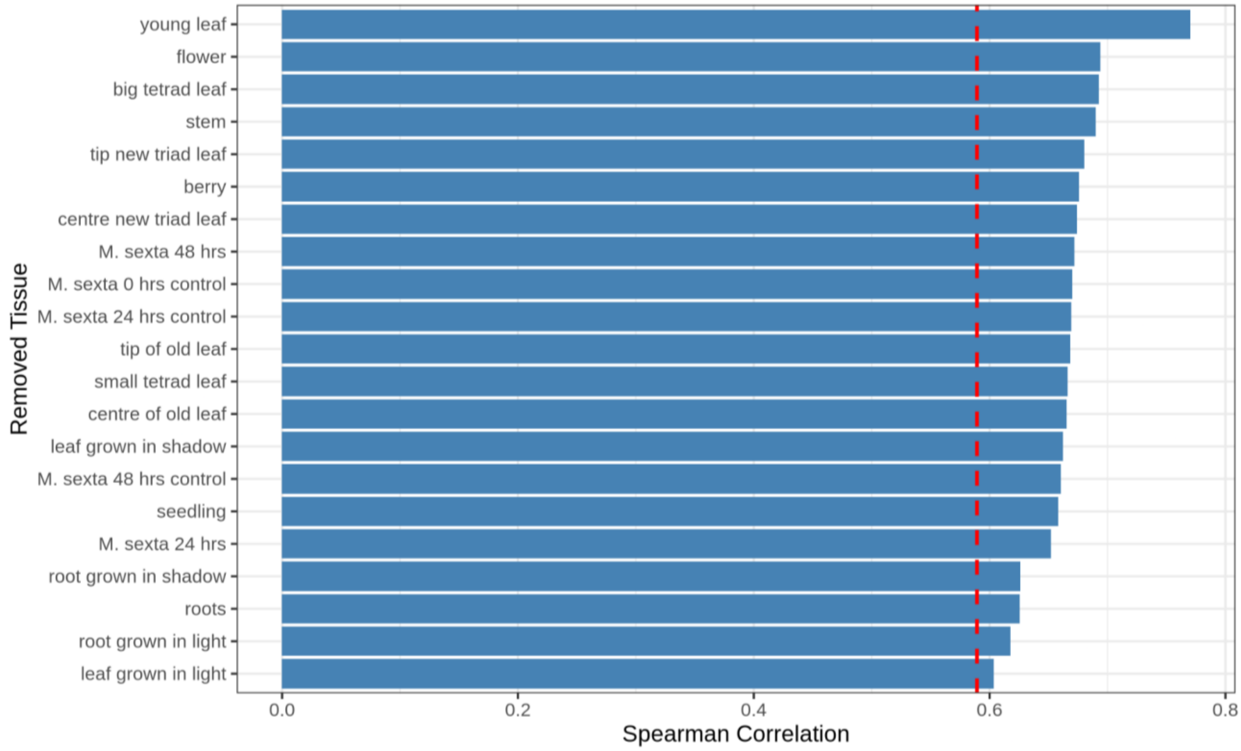


**Supplementary Figure S31: Leave-one-tissue-out (LOTO) correlation stability analysis between GS-11c and MSTRG.5531-11b homeolog across tissues:** Horizontal bar plot showing Spearman correlation coefficients between the GS-11c (Rte11cG086866.1) and MSTRG.5531-11b (Rte11bG086591.1) homeolog copies after iteratively excluding all samples from one tissue type at a time.

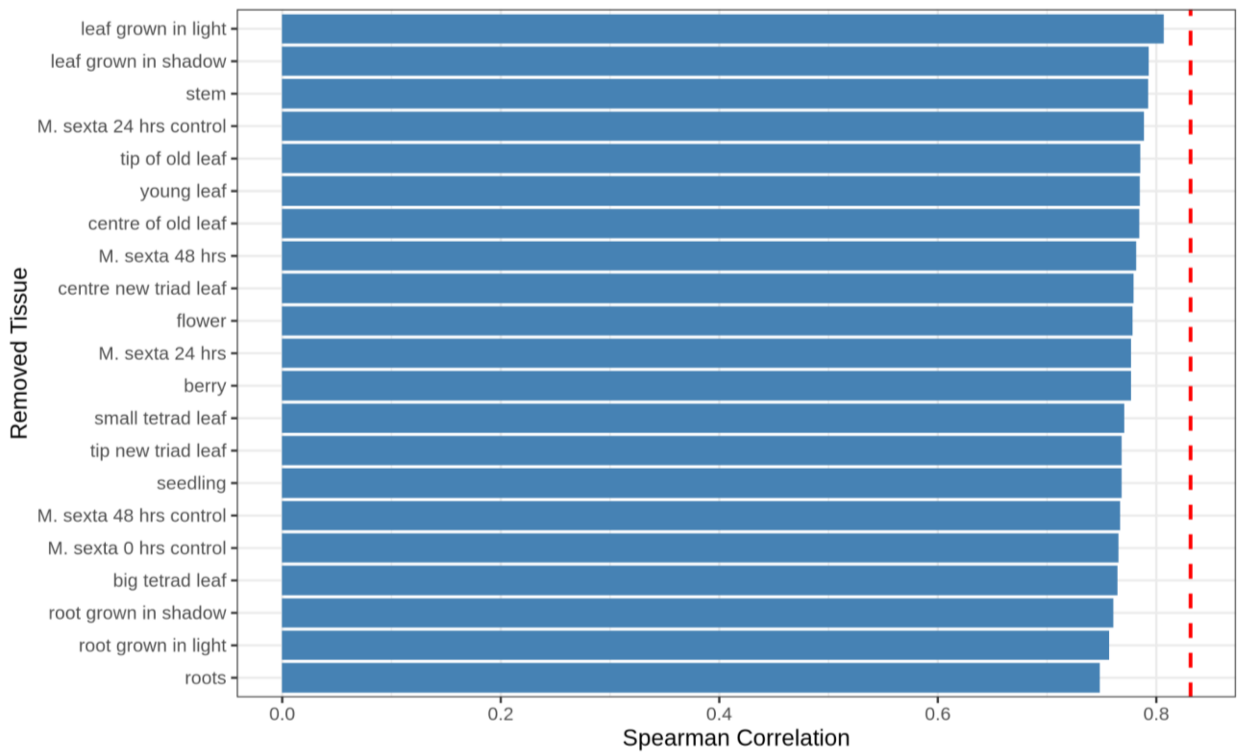


**Supplementary Figure S32: Leave-one-tissue-out (LOTO) correlation stability analysis between GS-11c and MSTRG.5531-11c homeolog across tissues:** Horizontal bar plot showing Spearman correlation coefficients between the GS-11c (Rte11cG086866.1) and MSTRG.5531-11c (Rte11cG086863.1) homeolog copies after iteratively excluding all samples from one tissue type at a time.

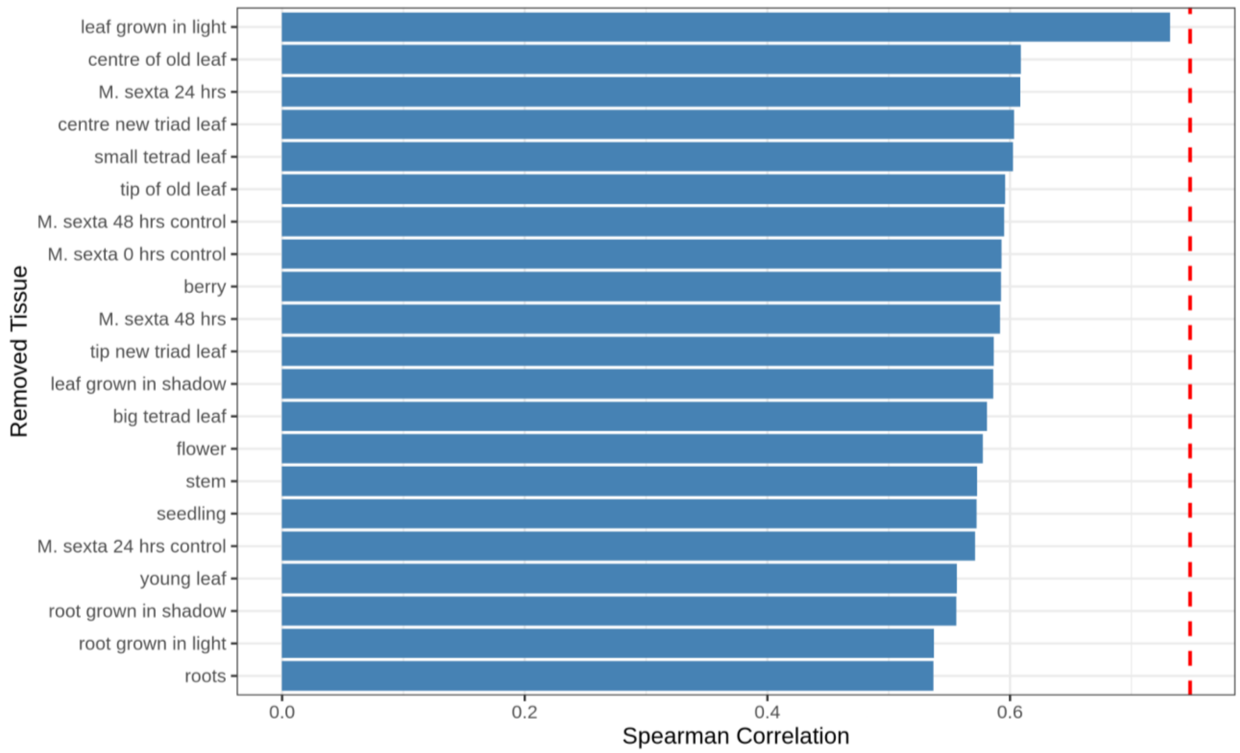


**Supplementary Figure S33: Leave-one-tissue-out (LOTO) correlation stability analysis between GS-11c and MSTRG.5530-11b homeolog across tissues:** Horizontal bar plot showing Spearman correlation coefficients between the GS-11c (Rte11cG086866.1) and MSTRG.5530-11b (Rte11bG086590.1) homeolog copies after iteratively excluding all samples from one tissue type at a time.
